# A novel hidden Markov approach to studying dynamic functional connectivity states in human neuroimaging

**DOI:** 10.1101/2022.02.02.478844

**Authors:** Sana Hussain, Jason Langley, Aaron R. Seitz, Xiaoping P. Hu, Megan A. K. Peters

## Abstract

**Introduction:** Hidden Markov models are a popular choice to extract and examine recurring patterns of activity or functional connectivity in neuroimaging data, both in terms of spatial patterns and their temporal progression. Although many diverse hidden Markov models have been applied to neuroimaging data, most have defined states based on activity levels (intensity-based states) rather than patterns of functional connectivity between brain areas (connectivity-based states), which is problematic if we want to understand connectivity dynamics: intensity-based states are unlikely to provide comprehensive information about dynamic connectivity patterns.

**Methods:** We addressed this problem by introducing a new hidden Markov model that defines states based on full functional connectivity profiles among brain regions. We empirically explored the behavior of this new model in comparison to existing approaches based on intensity-based or summed functional connectivity states using the HCP unrelated 100 functional magnetic resonance imaging “resting state” dataset.

**Results:** Our ‘full functional connectivity’ model discovered connectivity states with more distinguishable (i.e., unique and separable from each other) patterns than previous approaches, and recovered simulated connectivity-based states more faithfully than the other models tested.

**Discussion:** Thus, if our goal is to extract and interpret connectivity states in neuroimaging data, our new model outperforms previous methods which miss crucial information about the evolution of functional connectivity in the brain.

**Impact statement:** Hidden Markov models can be used to investigate brain states noninvasively. Previous models “recover” connectivity from intensity-based hidden states, or from connectivity ‘summed’ across nodes. Here we introduce a novel connectivity-based hidden Markov model and show how it can reveal true connectivity hidden states under minimal assumptions.

## 1. Introduction

The brain is a dynamical system of interacting and interchanging brain states (Chen et al., 2016; Lurie et al., 2020; Stevner et al., 2019; Vidaurre et al., 2017): patterns of activity levels or connectivity strengths that characterize and quantify network interactions. When extracted from functional magnetic resonance imaging (fMRI) data, these states can be categorized into two groups, defined either by activity levels of one or more nodes (brain areas; *intensity-based* states), or by patterns of functional connectivity between nodes (*connectivity-based* states). In contrast to intensity-based methods, connectivity-based states and their dynamics remain relatively underexplored. Developing and benchmarking new methods for extracting and characterizing these states is therefore of critical importance.

One promising approach used to characterize evolution of *intensity-based* brain states across time (i.e., not functional connectivity per se) is hidden Markov models (HMMs). HMMs utilize probabilistic methods to determine a hidden state sequence path not directly observable in data (Eddy, 1996, 2004; Jurafsky & Martin, 2009; L. R. Rabiner, 1989) by inferring underlying intensity-based states, where the probability of residing in any one of these states depends only on the previous state (Eddy, 1996, 2004; L. R. Rabiner, 1989). These models are powerful because (1) they have no assumptions about relationships among brain states (Chen et al., 2016), and (2) spatial and temporal information are inherent to the model. Because of these properties, HMMs have been used to identify latent brain states via signals acquired from fMRI and magnetoencephalography (Baker et al., 2014; Chen et al., 2016; Eavani et al., 2013; Stevner et al., 2019; Vidaurre, Abeysuriya, et al., 2018; Vidaurre et al., 2016, 2017; Vidaurre, Hunt, et al., 2018).

But what about connectivity-based states? Outside state-based analysis, a common approach is to calculate pairwise dynamic functional connectivity (dFC), i.e. the correlation in activity between pairs of nodes in a brain network and how such correlations change across time using a “sliding window” approach (Chen et al., 2016; Lurie et al., 2020; Vidaurre et al., 2017). However, to study connectivity-based states and their evolution, several groups have examined covariance values extracted from intensity-based HMMs, transforming them into Pearson correlations to create connectivity-like states (Chen et al., 2016; Stevner et al., 2019; Vidaurre, Abeysuriya, et al., 2018; Vidaurre et al., 2017). However, it is unclear to what extent such transformed covariances reflect true underlying *connectivity-based states* rather than simply the connectivities that the *intensity-based* states happened to exhibit. Other groups have used principal components analysis based approaches (Vidaurre, 2021; Vidaurre et al., 2021), or those that assume stationarity of intensity across time (Vidaurre, Abeysuriya, et al., 2018), with varying success. However, a more direct approach seems desirable – one which uses functional connectivity instead of signal intensity as a direct model input.

A more direct approach was recently undertaken by Ou and colleagues (2015), who summed the results from a dFC sliding window analysis into a representative “connectivity vector” -- describing a given node’s total connectivity to all other nodes in the network -- for every time point, and then fitted these with an HMM. Critically, however, this method sums over dynamic changes in pairwise connectivity, thereby potentially obscuring important changes in pairwise connectivities. For example, increased connectivity between the source node and one target node might be balanced by decreased connectivity to another node, such that no change is observed in overall connectivity. Importantly, interpretation of the output of a fitted HMM depends strongly on the inputs and assumptions used to develop the model, so the states resulting from Ou and colleagues’ (2015) method may differ from states recovered by a model fitted to *all* pairwise correlation values -- between *all* pairs of nodes -- within a sliding window.

Here we tackled these concerns by evaluating a new HMM-based method that fits all correlation values obtained from a dFC sliding window analysis. We comprehensively compared this novel *full functional connectivity* HMM (FFC HMM) to two previously-reported methods used to examine functional connectivity states in neuroimaging data: (1) a standard *intensity-based* HMM (IB HMM) (Chen et al., 2016; Stevner et al., 2019; Vidaurre et al., 2017), and (2) a *summed functional connectivity* HMM (SFC HMM) (Ou et al., 2015). We fitted each model to a widely-available existing dataset, the Human Connectome Project Unrelated 100 (Van Essen et al., 2013) resting state functional MRI dataset. Our findings highlight the advantages of our new FFC HMM in characterizing functional connectivity states, as well as cautioning against assuming that meaningful connectivity patterns can be derived from models fitted to alternative (intensity-based) or functional connectivity inputs summed across nodes.

## 2. Methods

### 2.1 HCP dataset and networks

All analyses were performed on the Human Connectome Project (HCP) Unrelated 100 (a subset of the S500 release) dataset (Van Essen et al., 2013) (https://ida.loni.usc.edu/login.jsp). We used 100 subjects (age = 22–36, gender = 54 female/46 male) who underwent a 14.4-minute resting state scan (repetition time = 720ms, flip angle = 52°, voxel size = 2mm^3^, echo time = 33ms, field of view = 208mm x 180mm). Data were preprocessed using the HCP minimal preprocessing pipeline: distortion correction, motion correction, alignment to standard space, and surface projection (Glasser et al., 2013). During development and fitting of the HMMs, one subject remained in one single, subject-specific state that was only visited by two other subjects for one timepoint each. Removing that subject did not affect the number of hidden states chosen (or any other parameter; data not shown), so we conducted all analyses on the remaining 99 subjects.

Following previous work (Deshpande et al., 2011; Raichle, 2011), BOLD signal was extracted from 29 regions of interest (ROIs) in four brain networks previously associated with resting state: the default mode network (DMN), fronto-parietal control network (FPCN), dorsal attention network (DAN), and salience network (SN). Nodes were defined using anatomical coordinates specified in literature (**Table A1**). See **Appendix A.1** for details.

### 2.2 Hidden Markov models

To evaluate the behavior of our novel *full functional connectivity* HMM (FFC HMM), we compared it to two previously-reported methods. The differences among these methods are defined by their inputs. Our FFC HMM takes as input a time series of pairwise dFCs between all pairs of ROIs. We compared this to models defined by: (1) BOLD time series (*intensity-based* HMM; IB HMM) (Chen et al., 2016; Stevner et al., 2019; Vidaurre et al., 2017), or (2) time series of dFC summed across all nodes that a given node is connected to (*summed functional connectivity* HMM; SFC HMM) (Ou et al., 2015). Each of these Gaussian HMMs relies on the same assumptions (Jurafsky & Martin, 2009; L. Rabiner & Juang, 1986; L. R. Rabiner, 1989). Models assumed the observation probability distribution is the normal distribution, and were implemented using the hmmlearn python library (Pedregosa et al., 2011) (**Appendix A.2**).

#### 2.2.1 Intensity-based HMM

The first comparison model is the standard *intensity-based* HMM (IB HMM) (Chen et al., 2016; Stevner et al., 2019; Vidaurre et al., 2017). BOLD signals from ROIs (**Table A1**) were extracted, preprocessed, z-scored, and concatenated across subjects (**Fig. A1**). These fMRI time series were concatenated timewise across all subjects to create a matrix of size (time * # subjects) x (# ROIs).

#### 2.2.2 Summed functional connectivity HMM

The second comparison model is the *summed functional connectivity* HMM (SFC HMM) (Ou et al., 2015). First, a sliding time window analysis (window length Δt=36 seconds) was used on the z-scored BOLD signal to obtain an ROI x ROI connectivity matrix of pairwise Pearson correlations between each ROI within each time window (Ou et al., 2015) (**Fig. A2**). This generated a connectivity time series of length (# TRs – Δt) representing the dynamics of functional connectivity over time. These square connectivity matrices were summed across one dimension to create a 1 x (# ROIs) vector depicting the total overall connectedness of each ROI to all other ROIs. Repeating this for every time window provided a “summed dFC time series” containing a (# time windows * # subjects) x ROI data matrix.

#### 2.2.3 Full functional connectivity HMM

To define our novel *full functional connectivity* HMM (FFC HMM), we aimed to remedy the shortcomings of SFC HMM. Our FFC HMM is therefore fitted to all correlation values in the lower (or, equivalently, upper) triangle of the dFC matrix in every time window, rather than summed connectivity vectors. Therefore, as before, a sliding window correlation analysis was performed on the z-scored BOLD signal with window length Δt = approximately 36 seconds (50 time points for primary analyses; see SFC HMM description), but instead of summing across one of the dimensions of the ROI x ROI matrices, the lower (or, equivalently, upper) triangle of R^2^ Pearson correlation values was restructured into a 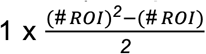 vector (**Fig. A3**). Repeating this for every time window gives a 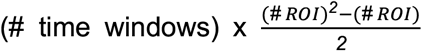 data matrix for every subject containing the time series of all pairwise connectivities between all pairs of ROIs. These results were then concatenated subject-wise as before to create a (# time windows * # subjects) x ROI data matrix.

### 2.3 Preliminary model fitting and analysis

We conducted several preliminary steps before proceeding to the full analyses. First, HMMs must be fitted with an *a priori* defined number of hidden states, so to determine the number of states for each model we adopted the Ranking and Averaging Independent Component Analysis by Reproducibility (RAICAR) method (Chen et al., 2016; Yang et al., 2008): briefly, this involved fitting the HMMs multiple times with different random seed initializations, aligning the recovered states by similarity, and seeking the number of states which produced maximal similarity across these aligned states (see **Appendix A.3** for details). Second, we quantified the degree to which each model tested could recognize ground truth in induced, or simulated, connectivity states (this approach follows similar logic to the RAICAR method and is described in detail in **Appendix A.4**). Finally, we evaluated the degree to which FFC HMM requires large amounts of data to produce robust results, by truncating the dataset and re-fitting the model (**Appendices A.5** & **C.2**). This is especially important to evaluate its utility for sample sizes smaller than the one used here. Intensity-defined state induction and model validation was also performed but is not included in the main text for brevity; see **Appendix B** for details.

### 2.4 Analysis of model outputs

#### 2.4.1 Connectivity state pattern analysis

Full connectivity state patterns (“connectivity state patterns” or “connectivity states”) depict the correlation strength between all pairs of nodes within a given state (i.e., each state consists of a 29×29 matrix of Pearson R^2^ values). The primary outcome metric of interest is the similarity across these recovered connectivity states from all three models. Due to model differences, the extraction method for connectivity states differed by model, but after extraction analyses were similar across models.

Both IB HMM and SFC HMM do not directly output connectivity profiles, so instead the full functional connectivity states must be recovered from model outputs. For IB HMM, connectivity states corresponding to each intensity state were acquired by mathematically transforming the covariance matrices outputted by the model fitting procedure into Pearson correlation values, as done previously (Eavani et al., 2013; Stevner et al., 2019; Vidaurre et al., 2017). (Note that this procedure demonstrates an assumption that such transformed covariance matrices may reveal connectivity states, whereas here we aimed to explicitly examine the relationship between transformed covariance based connectivity states and connectivity states inferred from models explicitly aimed to identify those states such as SFC HMM and FFC HMM.) For SFC HMM, the model’s outputs are vectors of mean summed correlation values representing global nodal strength during a time window which cannot be directly ‘unpacked’ into full pairwise connectivities among all nodes; therefore, we defined connectivity state patterns for SFC HMM by averaging connectivity matrices across time points when its Viterbi path labeled a state to be active. For FFC HMM, the connectivity state patterns are directly outputted from the model corresponding to the 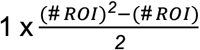 correlation vector inputted for every time window, which are then reformatted back into a symmetric ROI x ROI matrix to constitute the connectivity states.

All connectivity analyses were performed on *differential functional connectivity states*: connectivity matrices that “highlight” the unique functional connectivity characteristics of each state (Stevner et al., 2019). SFC and FFC HMM state differential functional connectivity states were computed using Eq. 1 where *X_i_* is the original raw functional connectivity matrix for state *i*, *H_i_* gives the differential functional connectivity states of *X_i_*, and *j* gives all state assignments (given the number of states determined for the model) excluding the value of *i* (Stevner et al., 2019).

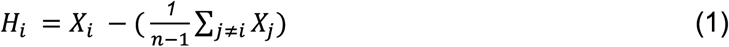

where *n* refers to the total number of states in the model. Here, the average is taken across the 7 states other than the current state being analyzed, so *n* − 1 = 7. Note that the values in the differential connectivity states represent connectivity levels relative to baseline, not the Pearson R^2^ values themselves: negative values are associated with below baseline correlations, not anticorrelations. Hereafter, “connectivity states” refers to differential connectivity states unless otherwise specified.

A critical question is to what extent FFC HMM’s recovered connectivity states could have been adequately captured by IB HMM and SFC HMM. Therefore, we examined the similarities among connectivity states recovered by IB, SFC, and FFC HMMs, with specific focus on SFC states versus FFC states. Connectivity state patterns were compared by Pearson correlation (Chen et al., 2016), similar to our procedure for the stability analysis (**Methods Section 2.3**, **Appendices A.3** and **C.1**). We sought to discover a ‘stability threshold’ (similar to the threshold of 0.9 used in the RAICAR analysis) that would result in unique, one-to-one pairwise matching between states recovered by two different models when states were aligned across those models by maximizing their similarity; see **Appendix A.4** for details of the model recovery process.

Finally, we also examined whether FFC HMM can recover summed connectivity vectors, as discovered by SFC HMM. Therefore, we computed the summed functional connectivity vectors corresponding to differential functional connectivity states for IB and FFC HMMs by summing the full 29×29 matrix of Pearson R^2^ values across one of the dimensions; for SFC, we utilized the summed connectivity vectors directly outputted by the model.

#### 2.4.2 Viterbi path analysis

We are also interested in examining trajectories through state space and how FFC HMM might differ from the two previous methods. The Viterbi path, or hidden state sequence, is directly outputted from all three HMMs. We examined a number of Viterbi path metrics, including switching rate, proportion of time spent in each state, the average duration of a state, and fractional occupancy correlation (Stevner et al., 2019; Vidaurre et al., 2017).

## 3. Results

### 3.1 Assessing model fits

The RAICAR-based stability analysis (**Methods Section 2.3**, **Appendix A.3**) determined that eight states were appropriate for all models (**Appendix C.1**).

To quantify the degree to which each model could recover ground truth in pure connectivity states, we also simulated connectivity-based “states” that were not accompanied by fluctuations in overall signal intensity (**Fig. 1a**), and used identical data preparation and fitting procedures to quantify how well each model could recover them (**Appendix A.4**). As expected, FFC HMM cleanly recovered simulated state trajectories (**Fig. 1d**), with mean correlation (R^2^) between true simulated and recovered Viterbi paths across subjects of 0.5738 ± 0.1301. Indeed, FFC HMM was more precise in its Viterbi path recovery than SFC HMM (**Fig. 1c**; mean R^2^ between simulated and recovered Viterbi paths of 0.3337 ± 0.1650). Unsurprisingly, IB HMM failed to adequately recover simulated pure connectivity states (**Fig. 1b**; mean R^2^ between simulated and recovered Viterbi paths of 0.1741 ± 0.1569), although it appeared to recognize when two networks were “turned on” in conjunction. This may have occurred because there was more connectivity for the model to recognize: both within- and between-network connectivity. However, recall that IB HMM does not natively evaluate or recover connectivity states, so connectivity must be estimated via the process described in **Methods Section 2.2.1**.

**Figure 1.**
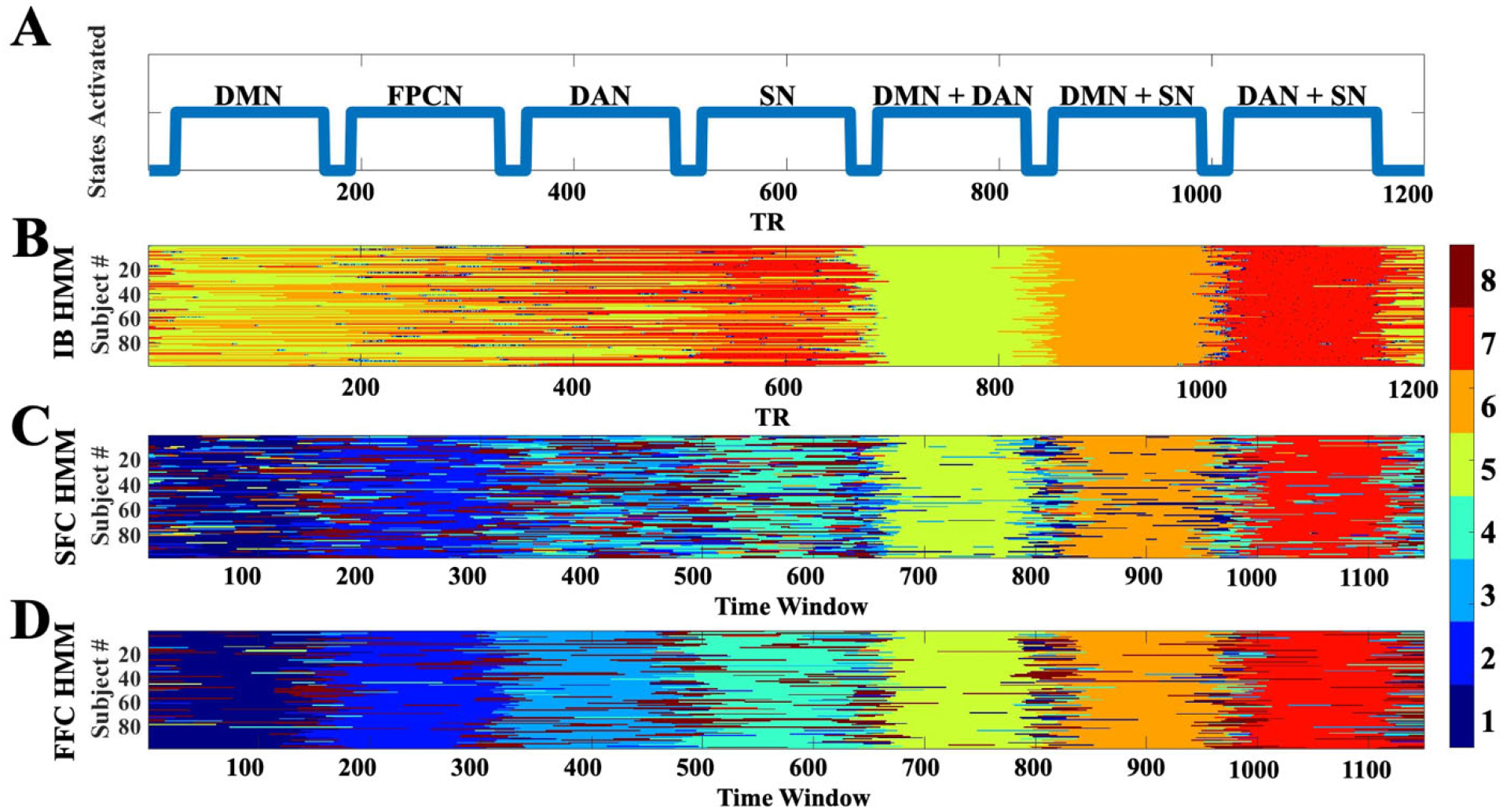
Verification of HMM connectivity-based states. **(A)** The artificially induced state sequence depicted which networks exhibited slightly increased within- and/or between-network connectivity. Outputted state sequences from **(B)** IB HMM, **(C)** SFC HMM, and **(D)** FFC HMM when connectivity states were induced. FFC HMM recovers simulated states better than the other two models; see main text for details.

FFC HMM also showed strong performance in recovering the connectivity states themselves. A paired t-test on the Fisher-z transformed R^2^ values between SFC and FFC HMMs indexing how similar each model’s recovered connectivity states were to the induced states showed that FFC HMM was significantly better at recovering induced states than SFC HMM (t(98) = 12.8745, p = 8.6087e-23). (See **Appendix B** for fuller discussion of the intensity-based states recovered by IB HMM.)

### 3.2 Analysis of model outputs

We next evaluated the states themselves as well as their Viterbi paths. States from each model are distinguished with subscripts corresponding to the HMM from which they stem, i.e., S1_FFC_ corresponds to State 1 from FFC HMM. We present results from IB and SFC HMMs first to provide context for the differences in behavior exhibited by FFC HMM.

#### 3.2.1 Connectivity state pattern analysis

First we examined IB HMM, which we did not expect to recover empirical connectivity states well. Unsurprisingly, the IB HMM differential functional connectivity states did not display strong distinguishing patterns either within or between the states (**Fig. 2**, top row). However, there were a few slight deviations from mean connectivity overall, especially in S4_IB_, S7_IB_, and S8_IB_. Nevertheless, these results show that a model not trained on connectivity states is, expectedly, not adept at recovering distinctive connectivity states.

**Figure 2.**
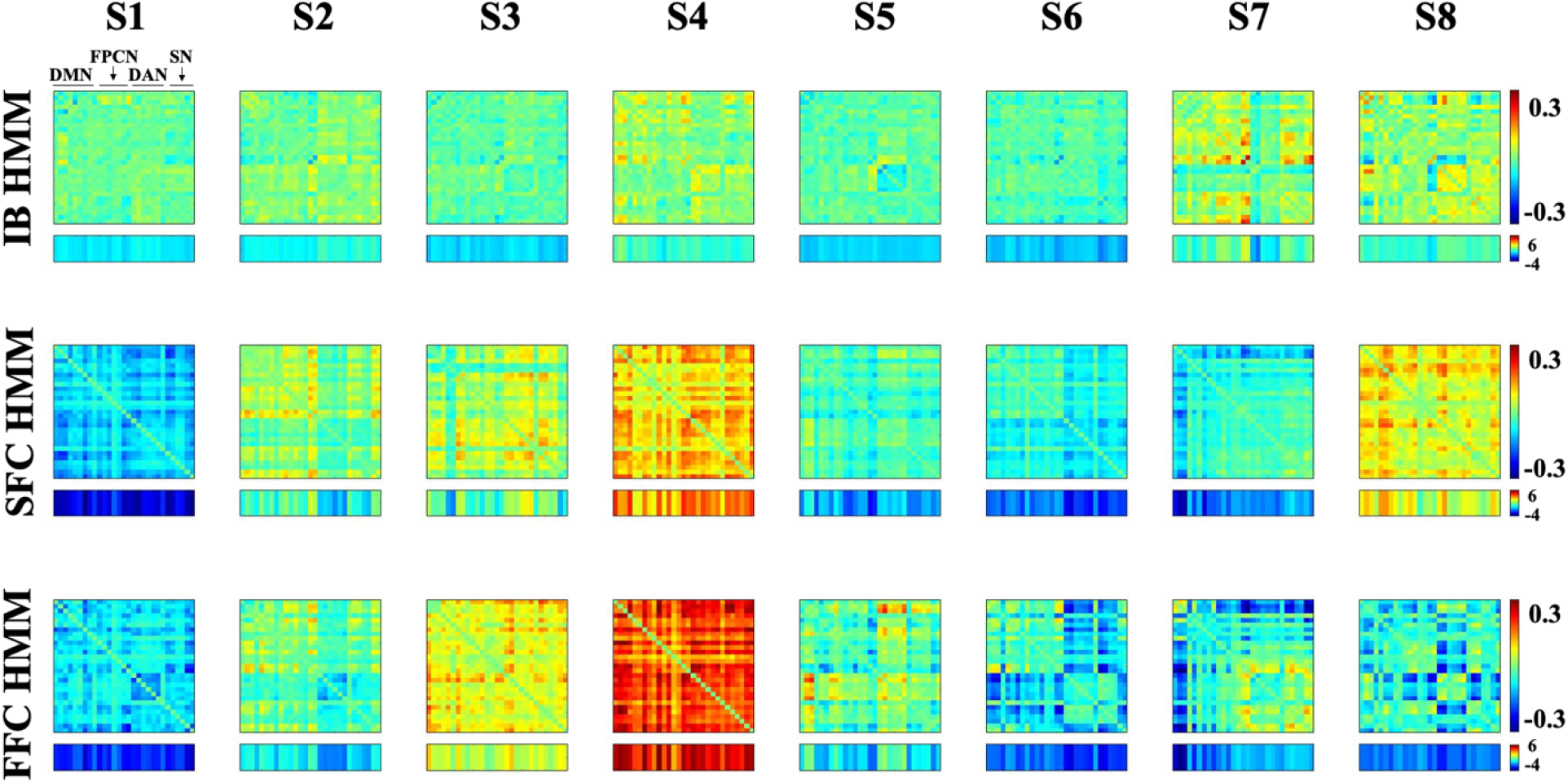
Differential functional connectivity states for SFC HMM (top row), FFC HMM (middle row), and IB HMM (bottom row). The summed connectivity vectors (summed across one dimension) are displayed below each state. The summed values for SFC were directly outputted from the model while those for IB and FFC were calculated as described in **Methods Section 2.4.1**.

The critical benchmark for FFC HMM is therefore SFC HMM’s behavior. Can a model fitted to summed connectivity vectors nevertheless adequately recover full connectivity profiles? We observed that SFC HMM extracted distinct differential functional connectivity profiles across all 8 states (**Fig. 2**, middle row). From visual examination, S1_SFC_ showed below-baseline correlations among all networks, while elevated correlations compared to baseline were seen overall in S4_SFC_ and S8_SFC_. Several states also showed distinct within-network changes in connectivity from baseline, with each state seeming to highlight a different network (e.g., S2_SFC_ excludes within-network connectivity in DAN, while S3_SFC_ and S5_SFC_ exclude within-network connectivity in DMN).

Crucially, to what extent are FFC HMM’s connectivity states different from SFC HMM’s? Visually, they appear to have poor correspondence (**Fig. 2**, bottom row). S4_FFC_ showed strong above-average connectivity across all nodes – far stronger than that shown in S4_SFC_ – and it was the only state to have overall above-average connectivity. In contrast, several other states showed a distinct reduction in functional connectivity specific to various networks. For example, S1_FFC_ shows a within-network DAN disconnect, while S5_FFC_ shows that DAN is connected to DMN and itself but not the other networks. Moreover, S6_FFC_ showed DAN and SN to be largely disconnected from DMN and FPCN, with DMN and FPCN connected to each other, while S7_FFC_ showed strong disconnection between DMN and all other networks. Other patterns can also be found. Importantly, though, no SFC HMM state exhibited any of these patterns.

We next engaged in a quantitative comparison between states recovered by each model in two ways. First, we computed the pairwise Pearson correlations among all pairs of states (**Fig. 3a**). A one-to-one match in states would be illustrated with one large correlation coefficient (one orange/yellow) square and seven small correlations (seven green/blue squares) in each row. Yet this phenomenon was not observed, as it appears there are several states that might equally ‘match’ across the two models from visual inspection.

**Figure 3.**
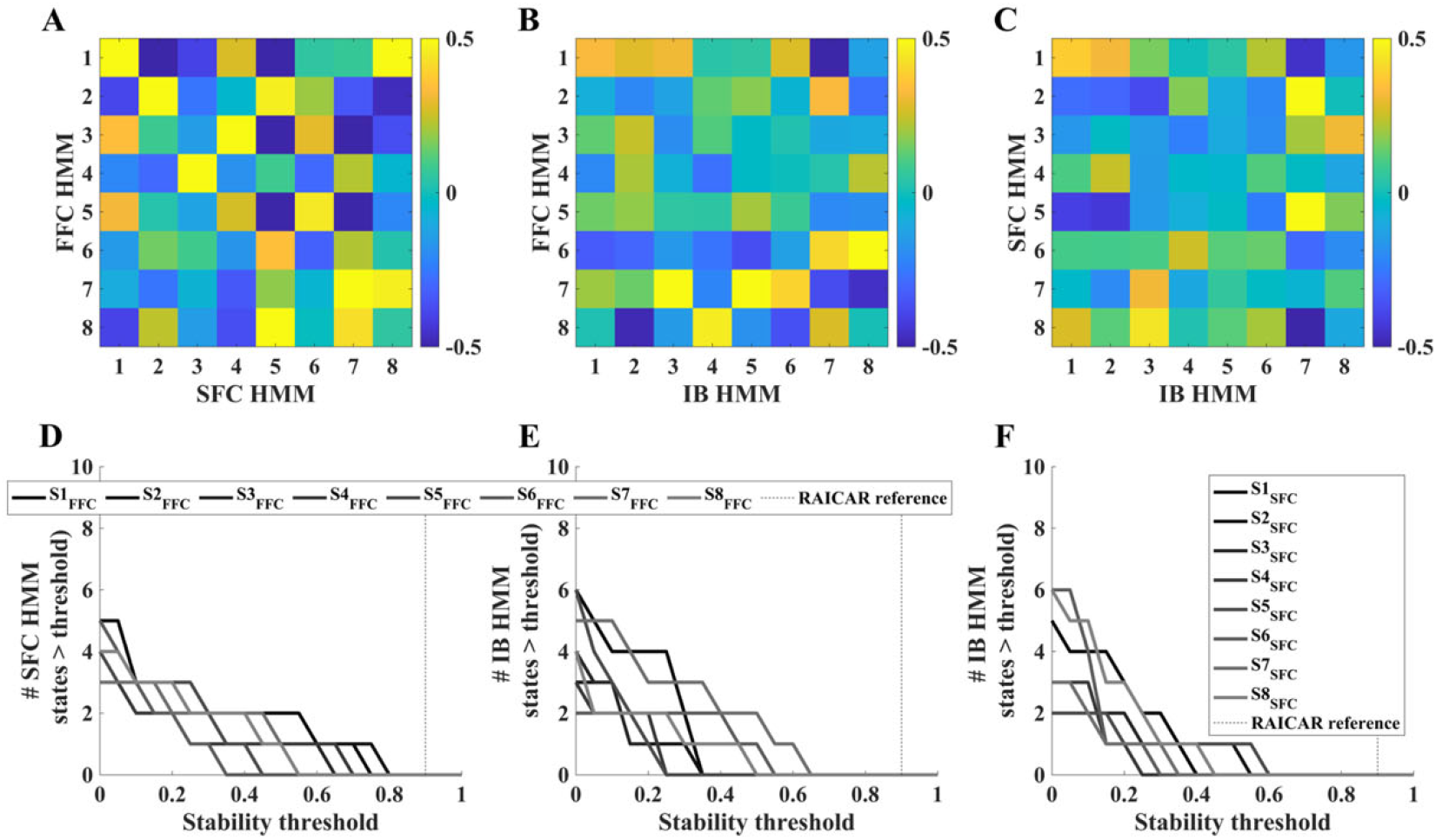
Pairwise comparisons between connectivity states discovered by each HMM show that each model recovered eight unique states. A one-to-one match between two states recovered by two different models would appear as a single orange/yellow square (high Pearson correlation) among seven green/blue squares (low Pearson correlation) in those two states’ row or column combination. However, **(A)** FFC HMM’s recovered states showed no unique correspondences based on similarity to those recovered by SFC HMM by visual inspection, and **(D)** no stability threshold (see the RAICAR analysis to discover number of hidden states, **Sections 2.3** & **3.1, Appendices A.3** & **C.1**) can lead to any semblance of a one-to-one match between FFC HMM’s recovered states and those recovered by SFC HMM. Similar results were found for pairwise comparisons between FFC HMM and IB HMM states (panels **B** and **E**) and between SFC HMM and IB HMM states (panels **C** and **F**).

Second, to quantitatively assess the degree to which there might be one-to-one state matching across states recovered by each pair of models, we took inspiration from the RAICAR analysis described above and sought to identify whether there was a threshold at which there would be a one-to-one state matching across all eight states found by both models. We are looking specifically for cases where a particular correlation threshold leads to exactly one SFC HMM state matching each of the FFC HMM states. We examined whether any possible correlation threshold in the range of 0-1 in steps of 0.05 would lead to exactly one SFC HMM state matching each FFC HMM state by counting the number of SFC states that exceeded each possible threshold. Visually, this accounts to counting how many ‘squares’ in each row of **Fig. 3a** surpass a particular value, with the goal being exactly one for all rows. This quantitative analysis confirmed the visual inspection results: There was no threshold for which all – or even most – FFC HMM states could achieve a unique mapping with exactly one SFC HMM state (**Fig. 3d**). This finding supports the interpretation that our novel FFC HMM recovers functional connectivity states that are distinct from those recovered by SFC HMM.

For completeness, we also repeated this analysis for comparisons between FFC HMM and IB HMM (**Figs. 3b** & **3e**), as well as comparing SFC HMM and IB HMM (**Figs. 3c** & **3f**). Results from these pairwise comparisons mirror those from the critical FFC vs SFC comparison: All three models appear to discover unique connectivity states, as there is no visual or quantitative correspondence between the states discovered by each model. Note that the comparisons with IB HMM are particularly informative, as they confirm that both FFC HMM and SFC HMM recovered states that reflected changes in connectivity patterns (in SFC HMM’s case, summed connectivity vectors) and were robust to fluctuations in overall BOLD signal magnitude (i.e., the connectivity-based models did not end up simply discovering ‘connectivity’ states based on fluctuations in average intensity). See **Appendix B** for fuller discussion of the intensity-based states recovered by IB HMM.

Finally, we also examined the relationship between summed functional connectivity vectors from FFC HMM and SFC HMM. We found that the mean absolute value of the inner product of the summed connectivity vectors for matched states from each model was very high (0.84), suggesting that FFC HMM’s outputs can be used to reconstruct what SFC HMM would have recovered.

#### 3.2.2 Viterbi path analysis

Viterbi paths can be visualized by assigning each state a color and plotting them for every person as a function of time point (TR in the fMRI time series, IB HMM, **Fig. 4a**) or time window (anchored on the first TR of the window, SFC HMM [**Fig. 4b**] and FFC HMM [**Fig. 4c**]) to show which states are active at each time point. These visualizations show that, compared to IB HMM, both connectivity-based HMMs appear ‘smoothed’ over time, i.e. transition among states more slowly; this is especially dramatic for FFC HMM. Autocorrelation and temporal discrepancy in input resolution between the intensity-versus connectivity-based HMMs -- i.e., IB HMM had a temporal resolution equal to that of the fMRI TR, while SFC and FFC HMMs had an effective sampling frequency on the order of 1 sample per 36 seconds -- likely contributing to this smoothing.

**Figure 4.**
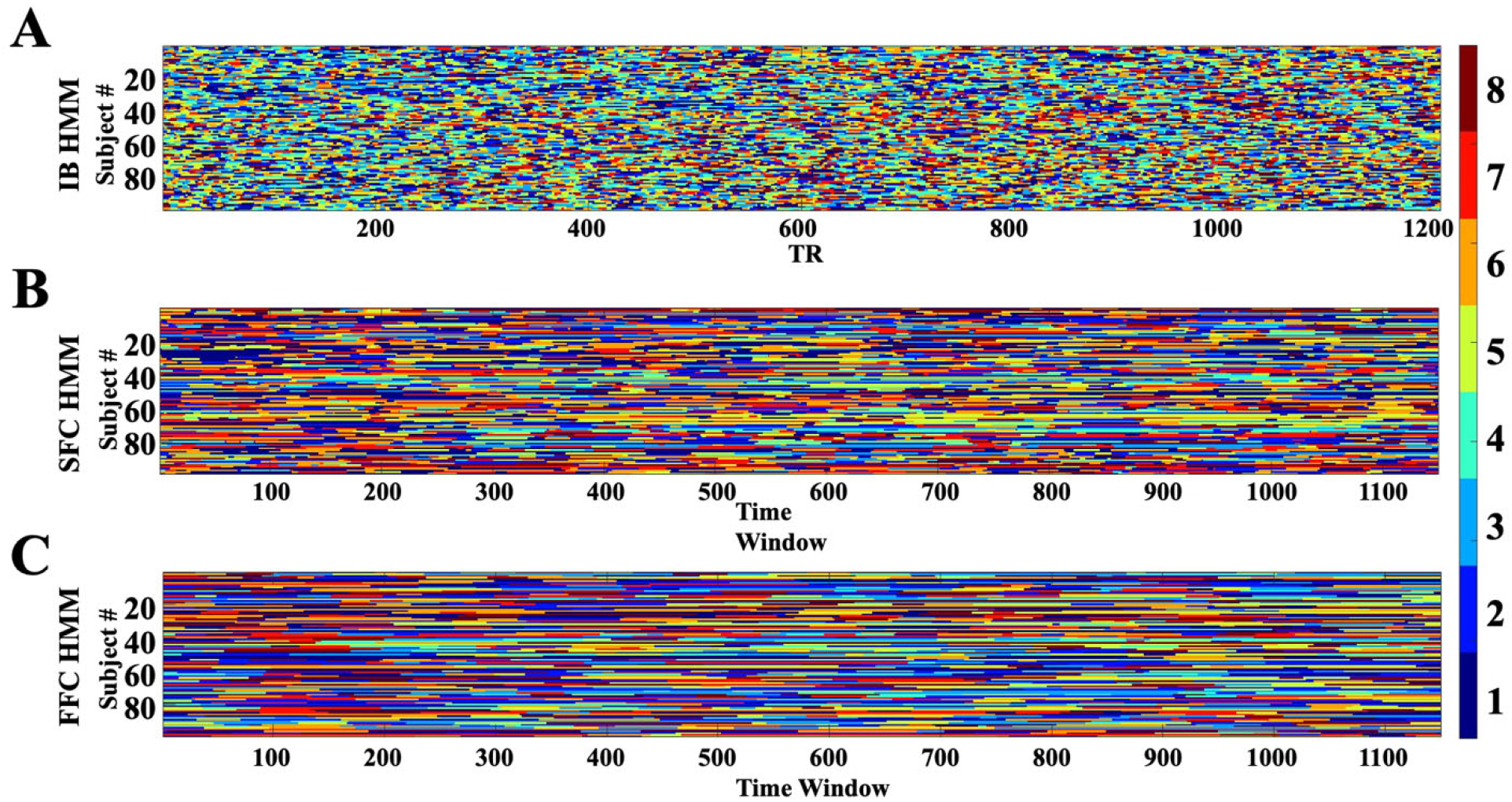
Viterbi Paths for **(A)** IB HMM, **(B)** SFC HMM, and **(C)** FFC HMM. The Viterbi paths for the SFC and FFC HMMs are much “smoother” (i.e., more spread out in time) than those of IB HMM. Within the connectivity-based HMMs, FFC HMM’s Viterbi path exhibits fewer and less frequent switches than SFC HMM’s, which may have occurred because of the selected number of hidden states or the total number of components fitted. See main text for detailed discussion.

Interestingly, this temporal ‘smoothing’ appears even more pronounced for FFC HMM than for SFC HMM – despite the fact that both SFC HMM and FFC HMM possess the same degree of ‘smoothing’ in the *input*, i.e. the same degree of autocorrelation induced by the sliding window computation of functional connectivity. We therefore suspect that FFC HMM’s complexity may have contributed to this behavior; we explore this possibility more in the **Discussion**.

As a final check, we also confirmed that the Viterbi paths were robust to different realizations, i.e. different initial conditions for model fitting. Recall that, although the RAICAR stability analyses determined 8 states was ideal for all three models, state assignment (e.g. labeling a state as “S1” vs “S8”) was initially arbitrary across all HMMs. Nevertheless, following state labeling alignment across initializations (see **Methods Section 2.3**, **Appendices A.3** and **C.1**), we observed that the Viterbi path was reproducible across different realizations of the HMMs (R^2^ ≥ ~0.84) for all models tested: For all initializations, all models recognized the same connectivity states at the same time windows, and the same switches between states.

## 4. Discussion

Here, we introduced a ‘full functional connectivity’ hidden Markov model (FFC HMM) and investigated its ability to extract functional connectivity states in resting state fMRI data. Using the HCP Unrelated 100, we fitted FFC HMM to a sliding window of functional connectivity from 29 ROIs across four networks (DMN, FPCN, DAN, and SN) and compared the connectivity states it discovered to those from two other models defined by summed functional connectivity (SFC HMM; (Ou et al., 2015)) and intensity (IB HMM; (Eavani et al., 2013; Stevner et al., 2019; Vidaurre et al., 2017))). FFC HMM’s full connectivity states were starkly different from those recovered by SFC and IB HMMs, even as FFC HMM recovered simulated connectivity-based states more faithfully than either IB or SFC HMM. Thus, we should not assume that connectivity states derived from intensity-based or summed functional connectivity based HMMs reflect true connectivity states; FFC HMM is a more appropriate choice to examine and interpret pure functional connectivity profiles when the question of interest is based on connectivity and not intensity fluctuations.

Interestingly, our results revealed that FFC HMM changed states more slowly even with the same sliding window size as SFC HMM. The slower switching rate in FFC HMM may be due to the increased number of components fitted in FFC HMM (406 connectivity values per time window) compared to SFC HMM (29 components); this may reduce state switching by requiring broader changes in connectivity across multiple nodes to register a change in functional connectivity state. Interestingly, this behavior may indicate that FFC HMM is less sensitive to noise than SFC HMM. Another possibility is that true functional connectivity may change quite slowly within a session, since it has been shown to be relatively stable across time with only moderate changes around the average (Vidaurre et al., 2021). Because SFC HMM sums across functional connectivities for a given node, such changes might accumulate and become comparably larger, causing SFC HMM to predict a state change when none is in fact present.

There are a number of key differences between our model and previous approaches that have targeted functional connectivity states using HMMs. In particular, Vidaurre and colleagues (Vidaurre, 2021; Vidaurre, Abeysuriya, et al., 2018; Vidaurre et al., 2021) fitted state-specific covariance matrices in a Gaussian HMM using intensity (BOLD or MEG-based) as inputs. Notably, their approach assumed that the mean intensity level of the observed data did not change between states, i.e. that mean activity was stationary across states (μ = 0 in the Gaussian model fitted by the HMM) thereby purposefully avoiding modeling changes in amplitude explicitly (Vidaurre, 2021). On the other hand, our approach uses covariance as the direct model input, sidestepping the assumption of stationarity in mean activity levels.

Other approaches have combined principal components analysis (PCA) and HMMs, either where PCA is done first and then the HMM fitted (Vidaurre et al., 2021), or where PCA and HMM fitting are done simultaneously (Vidaurre, 2021). In both cases, PCA is used to recover latent components in brain activity or connectivity data, which complicates the ability to interpret the discovered states with reference to known brain anatomy and functional networks. In the case of the second model (Vidaurre, 2021), this practice also suggests that the principal components discovered by the simultaneous approach can change as a function of connectivity state, such that the covariance matrix of one state may not refer to the same principal components as that of another state.

Finally, it has also been proposed that time-delay embedded HMMs may reveal functional connectivity networks in magnetoencephalography data (Vidaurre, Hunt, et al., 2018). Although this approach has the advantage of being purely data-driven and focused on phase-coupling network activity in the case of MEG data, it has not been applied to fMRI data and may encounter challenges due to its dependence on high temporal resolution and consequently long length of input data; we therefore leave exploration of this interesting possibility in fMRI data to future studies.

### 4.1 Limitations

HMMs assume independent Gaussians in the likelihood functions (Eddy, 1996, 2004; Jurafsky & Martin, 2009; L. Rabiner & Juang, 1986; L. R. Rabiner, 1989), but one might note that using sliding window functional correlations as inputs (SFC and FFC HMMs) introduces autocorrelation. While we assumed independence as a simplifying assumption (Ou et al., 2015), we also conducted an exploratory analysis which varied window size to explore potential impact of autocorrelation magnitude. This analysis revealed that even large changes in sliding window length do not meaningfully impact the connectivity states recovered by either SFC HMM or FFC HMM (**Appendix C.4**). Therefore, we can feel confident that autocorrelation concerns do not unduly influence our findings. In fact, we note that independence may not strictly be assumed even for IB HMM or similar models, as the hemodynamic response function is continuous and therefore autocorrelation from one TR to the next is likely due to the poor temporal resolution of fMRI in general. The lack of dependence on sliding window length may also speak to whether sliding windows in general are an appropriate choice for HMM-based analysis of resting state data, as this result may suggest that what the SFC and FFC HMMs may be recovering is a set of global, or relatively stable, functional connectivity states that the system slowly switches among as opposed to task-induced or more rapidly-changing dynamic states. Future research should investigate the relationship between global, unchanging dynamic connectivity profiles and the shifting profiles revealed by SFC HMM or FFC HMM, as well as in comparison to other methods proposed in the literature.

Gaussian HMMs also assume that the off-diagonal elements of the correlation matrices either are distributed normally, or that violations of this assumption do not unduly affect results (Chen et al., 2016). In the absence of a model which explicitly estimates the covariance of such off-diagonal elements, we evaluated the impact of Fisher-z transforming the Pearson correlations before using them as inputs so that they would occupy a range of [−∞,∞] rather than [−1,1]. Transforming the Pearson correlations did not significantly affect the connectivity states recovered by FFC HMM (**Appendix C.4**). We therefore elected to refrain from Fisher-z transforming to facilitate direct comparison to SFC HMM, which did not previously Fisher-z transform (Chen et al., 2016) and for which Fisher-z transformation is inappropriate because SFC HMM’s summed functional connectivity vectors already occupy a range of [−∞,∞].

Finally, our results are also limited by the sliding window approach, as a given brain state may not persist for the entire sliding window, or signals from multiple brain states may overlap in the window. Other approaches may appear to remedy this shortcoming, such as spatial independent component analysis (Beckmann et al., 2005; Smith et al., 2012), structural equation modeling (Schlösser et al., 2003), or coactivation patterns in intensity-based states (Liu et al., 2013; Liu & Duyn, 2013; Petridou et al., 2013). However, while these methods characterize spatial patterns of connectivity states, each time point is treated as independent and shuffling the time series does not affect spatial patterns of recovered brain states; they therefore only identify the states themselves and do not reveal the trajectory through state space. Thus, FFC HMM offers one of the best tools available to study the evolution of connectivity states over time.

### 4.2 Conclusions

We found that FFC HMM discovered connectivity states with more distinguishable patterns than those derived from HMMs with an intensity input (IB HMM) or summed connectivity input (SFC HMM), and which were fundamentally different from the functional connectivity profiles extracted by either of the other two methods. FFC HMM could also more faithfully recover simulated “ground truth” pure connectivity states. Because FFC HMM allows for a direct readout of connectivity-based states and their temporal evolution, it offers a powerful tool for extracting, analyzing, and understanding dynamic connectivities among brain regions.

## Data/Code Availability Statement

Data from HCP is available from the HCP database (https://ida.loni.usc.edu/login.jsp). Hidden Markov models were generated using the hmmlearn library in python (https://github.com/hmmlearn/hmmlearn).

## Credit Authorship Contribution Statement

<u>**Sana Hussain**:</u> Conceptualization, Formal Analysis, Investigation, Methodology, Project Administration, Software, Validation, Visualization, Writing -- Original Draft, Writing -- Review & Editing. <u>**Jason Langley**:</u> Investigation, Methodology, Supervision, Validation, Visualization, Writing -- Review & Editing. <u>**Aaron R. Seitz**:</u> Conceptualization, Funding Acquisition, Investigation, Methodology, Project Administration, Supervision, Validation, Visualization, Writing -- Review & Editing. <u>**Megan A. K. Peters**:</u> Conceptualization, Funding Acquisition, Investigation, Methodology, Project Administration, Resources, Supervision, Validation, Visualization, Writing -- Original Draft, Writing -- Review & Editing. <u>**Xiaoping P. Hu**:</u> Conceptualization, Funding Acquisition, Investigation, Methodology, Project Administration, Resources, Supervision, Validation, Visualization, Writing -- Review & Editing.

## Acknowledgements

This manuscript has been uploaded to the bioRxiv preprint server as Hussain, S., Langley, J., Seitz, A.R., Hu, X.P., & Peters, M.A.K. (2022). A novel hidden Markov approach to studying dynamic functional connectivity states in human neuroimaging. https://doi.org/10.1101/2022.02.02.478844.

## Appendices

## Appendix A. Additional methods details

All analyses presented in here used data from the HCP project (Principal Investigators: Bruce Rosen, M.D., Ph.D., Martinos Center at Massachusetts General Hospital; Arthur W. Toga, Ph.D., University of Southern California, Van J. Weeden, MD, Martinos Center at Massachusetts General Hospital). This project is supported by the National Institute of Dental and Craniofacial Research (NIDCR), the National Institute of Mental Health (NIMH) and the National Institute of Neurological Disorders and Stroke (NINDS). HCP is the result of efforts of co-investigators from the University of Southern California, Martinos Center for Biomedical Imaging at Massachusetts General Hospital (MGH), Washington University, and the University of Minnesota.

## A.1 Network ROI MNI coordinates

We narrowed our scope of analysis to four pre-defined networks that have previously been associated with resting state: the default mode network (DMN), fronto-parietal control network (FPCN), dorsal attention network (DAN), and salience network (SN). The nodes comprising each network were defined using anatomical coordinates specified in literature and converted from Talairach to MNI coordinates when necessary (Brett et al., 2002; Deshpande et al., 2008, 2009; Laird et al., 2005; Lancaster et al., 2007; Raichle, 2011; Stilla et al., 2007). Dorsal anterior cingulate cortex and left dorsolateral prefrontal cortex ROIs in FPCN and right parahippocampal gyrus and right inferolateral temporal cortex in DMN were excluded from analyses because they overlapped or were too closely located to other ROIs. Thus, we only used 29 ROIs per subject when examining the HCP dataset (9 from DMN, 7 from FPCN, 6 from DAN, and 7 from SN).

Talairach coordinates for DMN, FPCN, and DAN were taken from Deshpande et al. 2011 and converted to Montreal Neurological Institute (MNI) coordinates (Brett et al., 2002; Deshpande et al., 2011; Laird et al., 2005; Lancaster et al., 2007). MNI coordinates for SN were taken directly from Raichle (2011) 2011 (Raichle 2011). MNI coordinates for all ROIs can be found in **Table A1**. After labeling each ROI with a 5mm^3^ isotropic marker, the BOLD signal was extracted from each voxel and averaged across all voxels in the ROI, producing a single time series representing the behavior of the ROI as a whole. This procedure was repeated for every ROI in a network for a total of 29 ROIs (9 from DMN, 7 from FPCN, 6 from DAN, and 7 from SN) per subject (Deshpande et al., 2008, 2009; Stilla et al., 2007).

**Table A1:** List of MNI coordinates used for ROIs in the default mode network (DMN), fronto-parietal control network (FPCN), dorsal attention network (DAN), and salience network (SN). Talaraich coordinates for DMN, FPCN, and DAN were taken from Deshpande et al. (2011) and were converted to MNI using methods described previously (Brett et al., 2002; Deshpande et al., 2011; Laird et al., 2005; Lancaster et al., 2007) while MNI coordinates for SN were taken directly from Raichle (2011).

| ROI | Abbrev. | Full Name | MNI (x,y,z) | Source |
| --- | --- | --- | --- | --- |
| **Default Mode Network** |  |  |  |  |
| 1 | PCC | Posterior Cingulate Cortex | (2,54,16) | Deshpande et al. 2011 |
| 2 | L pIPL | Left Posterior Inferior Parietal Lobule | (-46,-72,28) | Deshpande et al. 2011 |
| 3 | R pIPL | Right Posterior Inferior Parietal Lobule | (50,-64,26) | Deshpande et al. 2011 |
| 4 | OFC/vACC | Orbitofrontal Cortex/Ventral Anterior Cingulate Cortex | (4,30,26) | Deshpande et al. 2011 |
| 5 | dMPFC BA 8 | Dorsomedial Prefrontal Cortex Brodmann Area 8 | (-14,54,34) | Deshpande et al. 2011 |
| 6 | dMPFC BA 9 | Dorsomedial Prefrontal Cortex Brodmann Area 9 | (22,58,26) | Deshpande et al. 2011 |
| 7 | L DLPFC | Left Dorsolateral Prefrontal Cortex | (-50,20,34) | Deshpande et al. 2011 |
| 8 | L PHG | Left Parahippocampal Gyrus | (-10,-38,-2) | Deshpande et al. 2011 |
| 9 | L ITC | Left Inferolateral Temporal Cortex | (-60,-20,-18) | Deshpande et al. 2011 |
| **Fronto-Parietal Control Network** |  |  |  |  |
| 1 | L aPFC | Left Anterior Prefrontal Cortex | (-36,56,10,) | Deshpande et al. 2011 |
| 2 | R aPFC | Right Anterior Prefrontal Cortex | (34,52,10) | Deshpande et al. 2011 |
| 3 | R DLPFC | Right Dorsolateral Prefrontal Cortex | (46,14,42) | Deshpande et al. 2011 |
| 4 | L aINS | Left Anterior Insula | (-30,20,-2) | Deshpande et al. 2011 |
| 5 | R aINS | Right Anterior Insula | (32,22,-2) | Deshpande et al. 2011 |
| 6 | L aIPL | Left Anterior Inferior Parietal Lobule | (-52,-50,46) | Deshpande et al. 2011 |
| 7 | R aIPL | Right Anterior Inferior Parietal Lobule | (52,-46,46) | Deshpande et al. 2011 |
| **Dorsal Attention Network** |  |  |  |  |
| 1 | L MT | Left MidThalamus | (-44,-64,-2) | Deshpande et al. 2011 |
| 2 | R MT | Right MidThalamus | (50,-70,-4) | Deshpande et al. 2011 |
| 3 | L FEF | Left Frontal Eye Field | (-24,-8,50) | Deshpande et al. 2011 |
| 4 | R FEF | Right Frontal Eye Field | (28,-10,50) | Deshpande et al. 2011 |
| 5 | L SPL | Left Superior Parietal Lobule | (-26,-52,56) | Deshpande et al. 2011 |
| 6 | R SPL | Right Superior Parietal Lobule | (24,-56,-54) | Deshpande et al. 2011 |
| **Saliience Network** |  |  |  |  |
| 1 | DAC | Dorsal Anterior Cingulate | (0,-22,36) | Raichle 2011 |
| 2 | L aPFC | Left Anterior PFC | (-34,44,30) | Raichle 2011 |
| 3 | R aPFC | Right Anterior PFC | (32,44,30) | Raichle 2011 |
| 4 | L Insula | Left Insula | (-40,2,6) | Raichle 2011 |
| 5 | R Insula | Right Insula | (42,2,6) | Raichle 2011 |
| 6 | L LP | Left Lateral Parietal | (-62,-46,30) | Raichle 2011 |
| 7 | R LP | Right Lateral Parietal | (62,-46,30) | Raichle 2011 |

## A.2 HMM models: details & graphical descriptions

All models were Gaussian hidden Markov models (that is, they assume the observation probability distribution is the normal distribution), fitted with the hmmlearn package using standard procedures (Pedregosa et al., 2011). The forward and Viterbi algorithms were used in conjunction to identify the most likely sequence of hidden states given the observable BOLD signal. The Baum-Welch algorithm was then implemented to calculate the transition and emission probabilities of a given state (Jurafsky & Martin, 2009; L. Rabiner & Juang, 1986; L. R. Rabiner, 1989).

The following figures graphically describe the methods used to implement the three HMMs discussed in the main text. **Fig. A1** corresponds to IB HMM, **Fig. A2** to SFC HMM, and **Fig. A3** to FFC HMM.

## A.2.1 Intensity-based HMM

**Figure A1.**
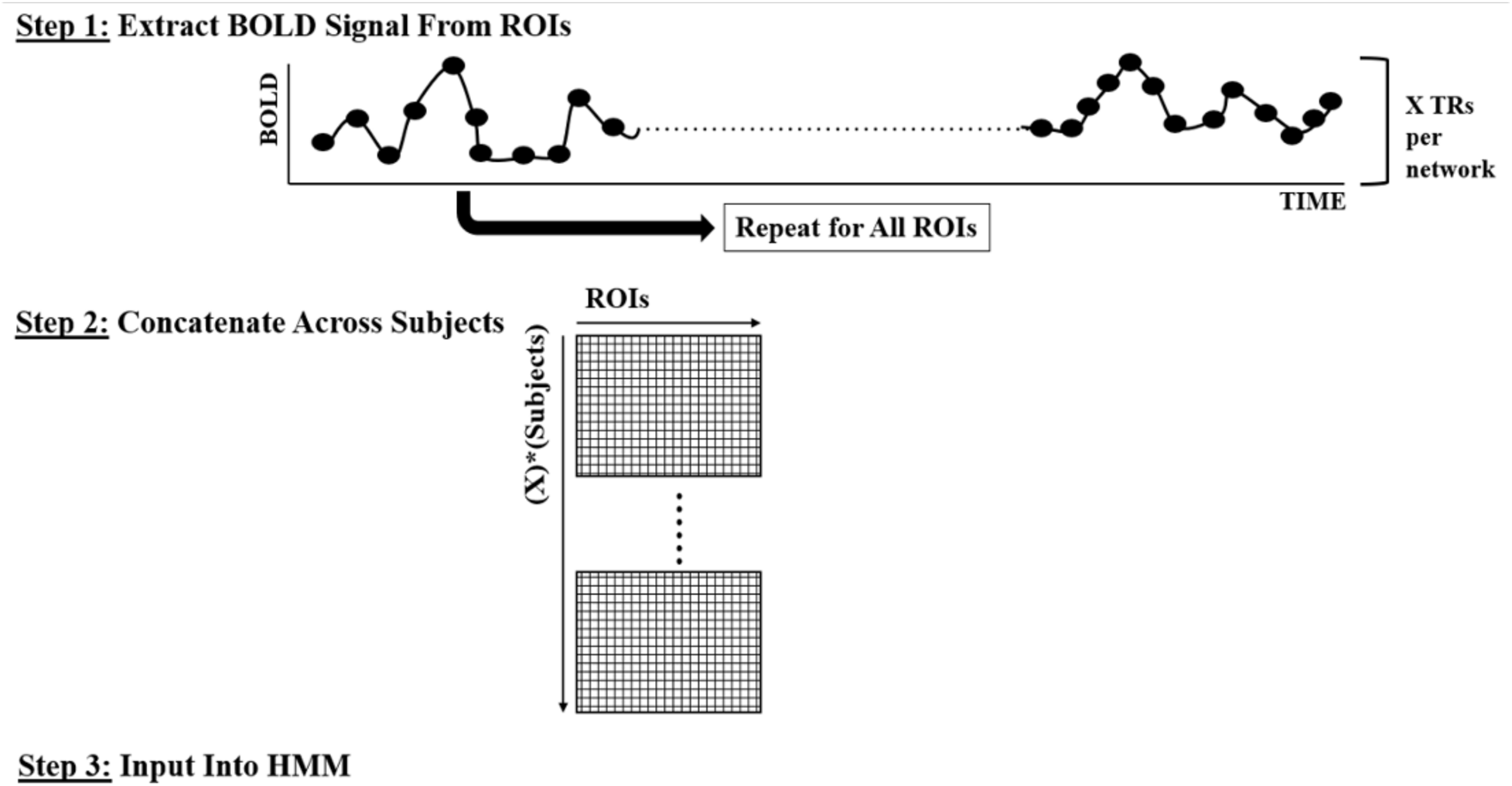
Illustration of the procedure used to implement IB HMM. Once the BOLD signal is extracted from all predefined ROIs, it is concatenated across subjects and fitted with an HMM from the python hmmlearn library (Pedregosa et al., 2011).

## A.2.2 Summed functional connectivity HMM

**Figure A2.**
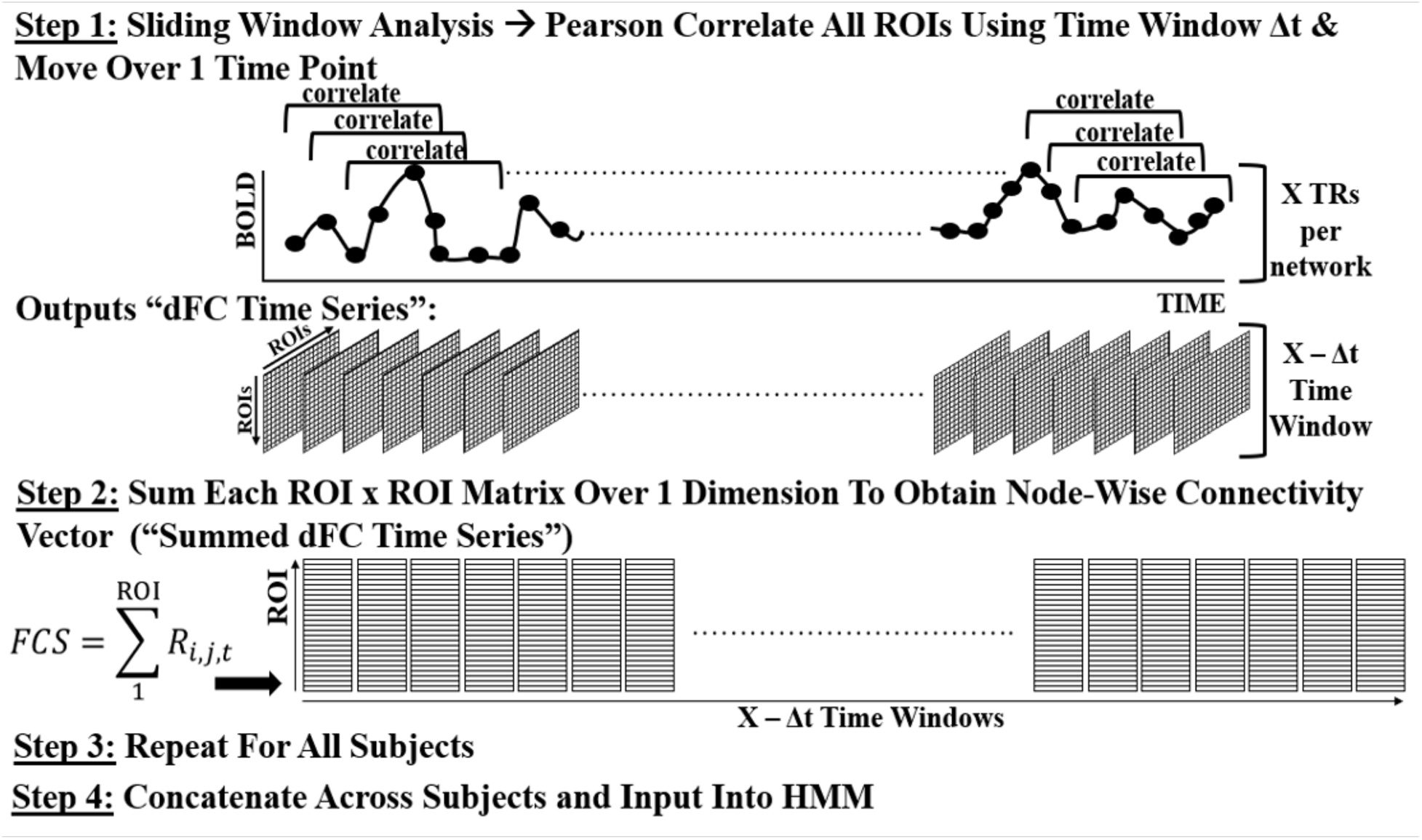
Illustration of the procedure used to implement SFC HMM as followed by the method described in Ou et al. (2015). Once a sliding time window correlation analysis is performed, the connectivity matrix in each time window is summed across a dimension into a representative nodal connectivity vector. After executing this for all time windows and for all subjects, the data is concatenated across all subjects and fitted with an HMM from the python hmmlearn library (Pedregosa et al., 2011).

## A.2.3 Full functional connectivity HMM

**Figure A3.**
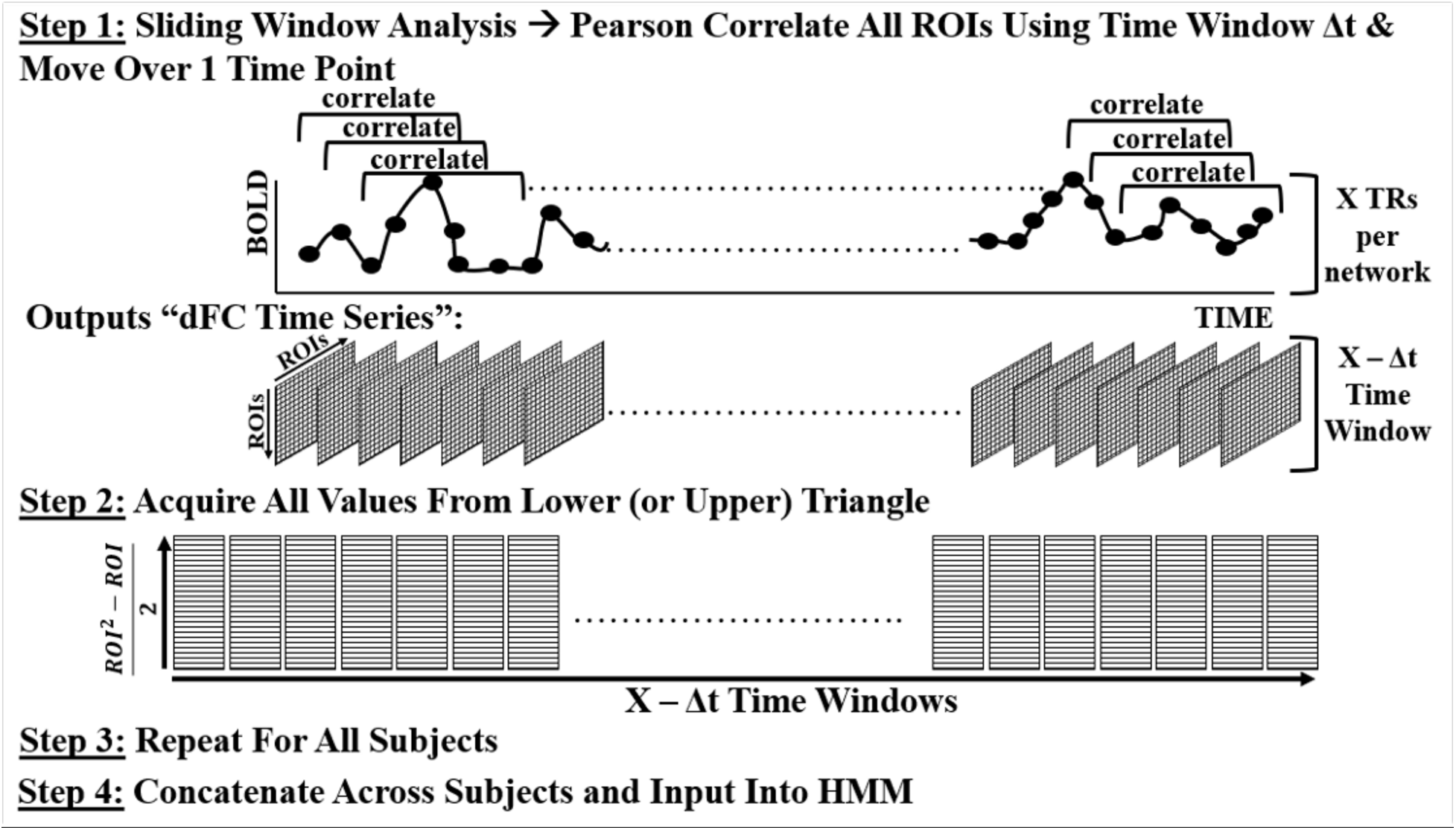
Illustration of the procedure used to implement FFC HMM. Once a sliding time window correlation analysis is performed, the lower (or upper) triangle of values from the connectivity matrix in each time window is flattened into a vector. After executing this for all time windows and for all subjects, the data is concatenated across all subjects and fitted with an HMM from the python hmmlearn library (Pedregosa et al., 2011).

## A.3 Detail of the RAICAR-based method to determine number of hidden states for each model

We examined the stability of models with 3 to 15 hidden states. For each HMM with each number of hidden states, three sets of state patterns were obtained by three different realizations: one with uniform starting probability of residing in all states, and two with randomly assigned starting probabilities. Because the labeling of states is arbitrary, there is no reason to believe State 1 will match across realizations even if a given number of hidden states is optimally stable. Therefore, state patterns were matched across realizations via Pearson correlations, such that e.g. State 1 from Realization 2 was relabeled as State 2 just in case the Pearson correlation between that state and State 2 from Realization 1 was higher than any other pairwise correlation. Thus, after relabeling, each state label across realizations universally corresponded to the same spatial pattern to the maximal extent possible within stability constraints. Within each state assignment, the matched state patterns were then Pearson correlated between all realizations to obtain (*# states*)!/*2*! * (*#states* − *2*)! Pearson’s R^2^ values, thereby determining the maximal degree of similarity between the matched patterns for the present number of hidden states being tested. Finally, these values were then averaged, sorted from largest to smallest, and plotted as a function of the number of states in the model.

We repeated this process for a range of number of hidden states, generating a dataset of pattern similarity (R^2^) organized by pattern label; these maximal achievable pattern similarities were compared against a predetermined threshold of stability of 0.9. Previous groups that used the RAICAR-based method examined more ROIs (236 from Chen et al. (2016) and 162 independent components from Yang et al. (2008)) and employed a stability threshold of 0.8 (Chen et al. 2016; Yang et al. 2008). As we explored only 29 ROIs, we opted to appoint a more conservative threshold: 0.9. Cases where R^2^ values began to dip below this threshold indicated that this model was unstable with that number of hidden states, because the states were not matching sufficiently (Chen et al., 2016; Yang et al., 2008).

**Fig. A4** illustrates the procedure used to determine the optimal number of states for this investigation: a Ranking and Averaging Independent Component Analysis by Reproducibility (RAICAR) based method.

**Figure A4.**
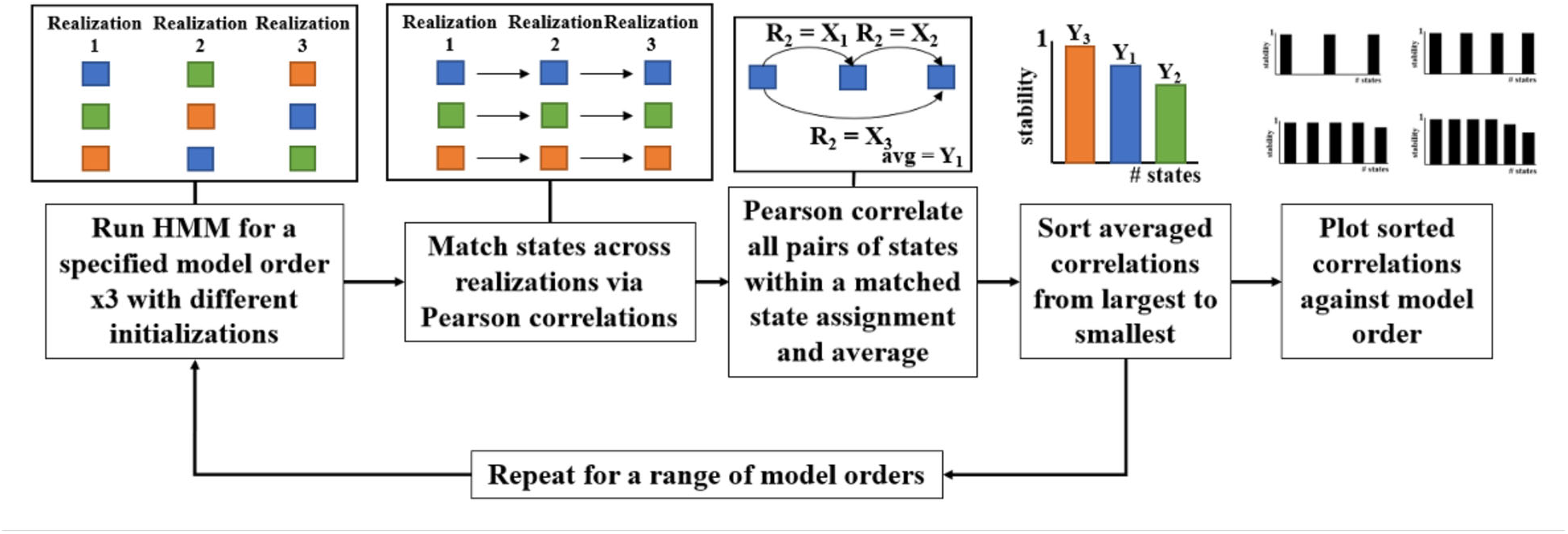
Schematic illustrating the procedure for the RAICAR-based method for determining number of hidden states in a model. Each of the blocks shown represent a state where matched states are shown in the same color. A basic example of 3 states is shown where states’ spatial patterns are first matched across realizations using Pearson correlation values (Chen et al., 2016; Yang et al., 2008). That is, states’ spatial patterns have been matched via the highest R^2^ value observed and reordered to the same state assignment. The Pearson correlation is then found amongst all pairs of states within a matched group and averaged to represent the stability of that state pattern for that number of hidden states. This procedure is then repeated for all states, with a model fitted with that particular number of hidden states.

## A.4 Details of model recovery

To validate our FFC HMM and benchmark it against previous approaches, we verified the degree to which it can recognize ground truth by successfully recovering predefined pure connectivity states that were not accompanied by fluctuations in mean activation level. The purpose of this analysis was to evaluate the degree to which each HMM could recover states defined purely by fluctuations in connectivity, i.e. which were not at all related to fluctuations in overall activity level. Thus, we manipulated the HCP data to induce “connectivity-defined states” and evaluated its performance as follows. First, the preprocessed data extracted from our 29 ROIs were randomly permuted across time to create a noisy time series. To create “connectivity-defined” ground truth states (absent systematic fluctuations in intensity), we replaced the scrambled data from these same time points with values drawn from a multivariate gaussian distribution using μ = 0 and an ROI x ROI σ matrix consisting of 0.1 on the off-diagonals where we aimed to induce connectivity, and ones on the diagonals. For example, to induce a “DMN (9 ROIs) connectivity state”, a 9 x 9 σ matrix was made with 0.1 on all off diagonals and 1 on the diagonals, which was then used to seed a time series of simulated BOLD response within the time points for which this connectivity state was to be induced; the result was then used to replace the temporally scrambled data. Connectivity state patterns and a hidden state sequence were acquired from FFC HMM outputs and compared against outputs from IB HMM and SFC HMM to establish to what extent each model is capable of recovering ground truth in pure connectivity-defined states.

## A.5 Details of model robustness to data size

To determine the ability of each model to recover stable states regardless of the length of the resting state scan, we performed an additional reproducibility analysis based on truncated datasets. To evaluate whether each model could stably discover the same states using a significantly reduced dataset, we fit the IB, SFC, and FFC HMMs to only the first half of the resting state scan for all subjects (that is, the first 7.2 minutes of a 14.4 minute scan) and computed the similarity between the discovered states for this ‘half’ dataset and the full dataset used in the main analyses. Similarity was computed according to the same process used in the RAICAR-based stability analysis (**Methods Section 2.3**, **Appendix A.3**), except that states were matched between the half- and full-dataset HMMs (fit with the uniform initial conditions) rather than between different realizations.

In **Fig. A5**, we present the connectivity-based state patterns identified (along with their corresponding summed vectors) when only half of the fMRI dataset (7.2 minutes) was fitted with an HMM.

**Figure A5.**
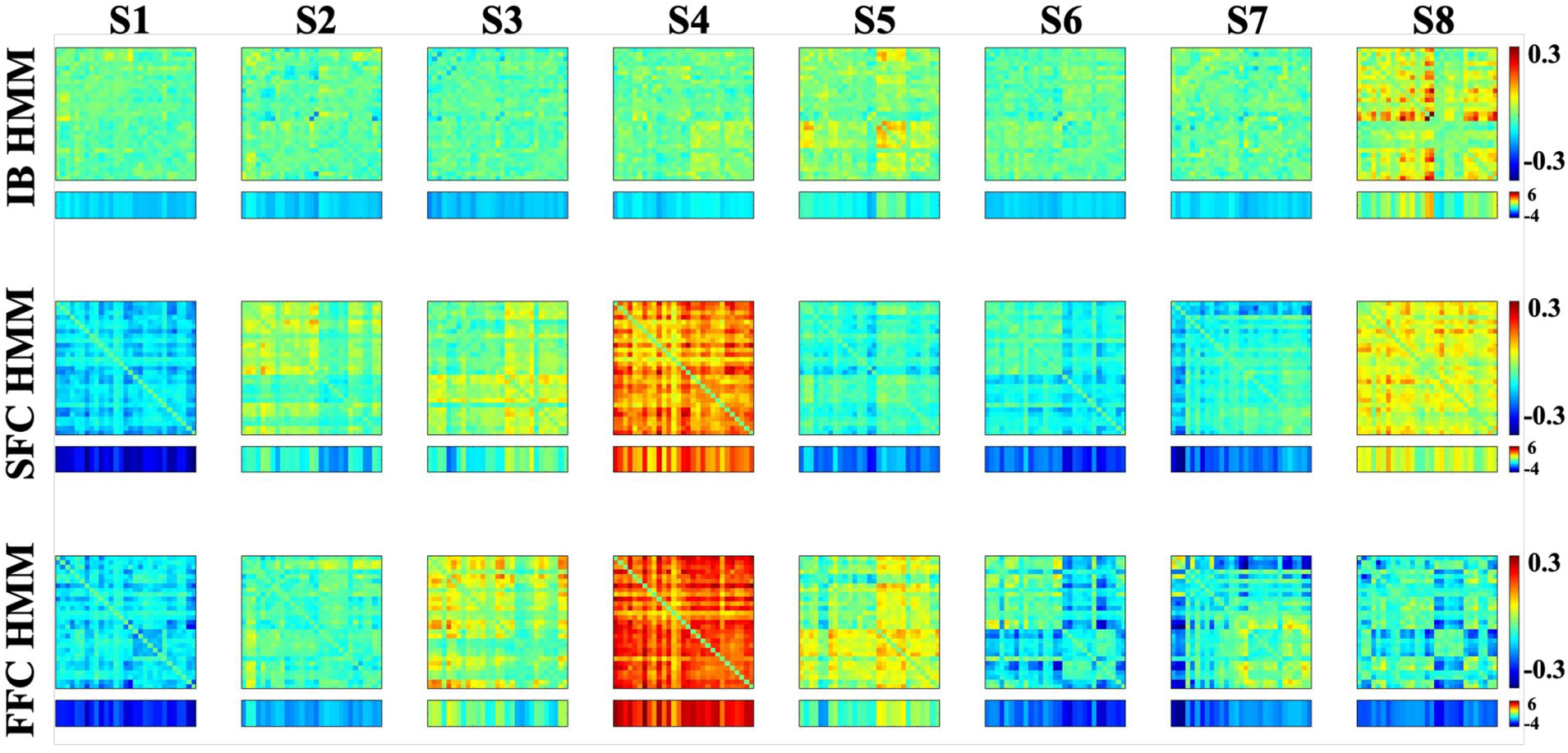
Differential functional connectivity states for SFC HMM (top row), FFC HMM (middle row), and IB HMM (bottom row) when only half of the fMRI dataset (7.2 minutes) were used. The summed connectivity values across one dimension are displayed below each state.

## Appendix B. Intensity-based state patterns

## B.1 Pattern acquisition

Intensity state patterns are defined as a combination of activated or deactivated ROIs comprising the four aforementioned networks (i.e., each state consists of a 1×29 vector of intensity levels). Intensity states in the SFC and FFC HMMs were computed using the method of state acquisition described by Chen and colleagues (Chen et al., 2016). **Fig. B1** shows that the BOLD time series was truncated by a length of Δt/2 to match the temporal scale of the SFC and FFC Viterbi paths. Next, the BOLD signal at every TR where the SFC or FFC labeled a state to be active was averaged. Repeating this for all ROIs yielded a 1 X ROI vector of intensity for every ROI for a particular state.

**Figure B1.**
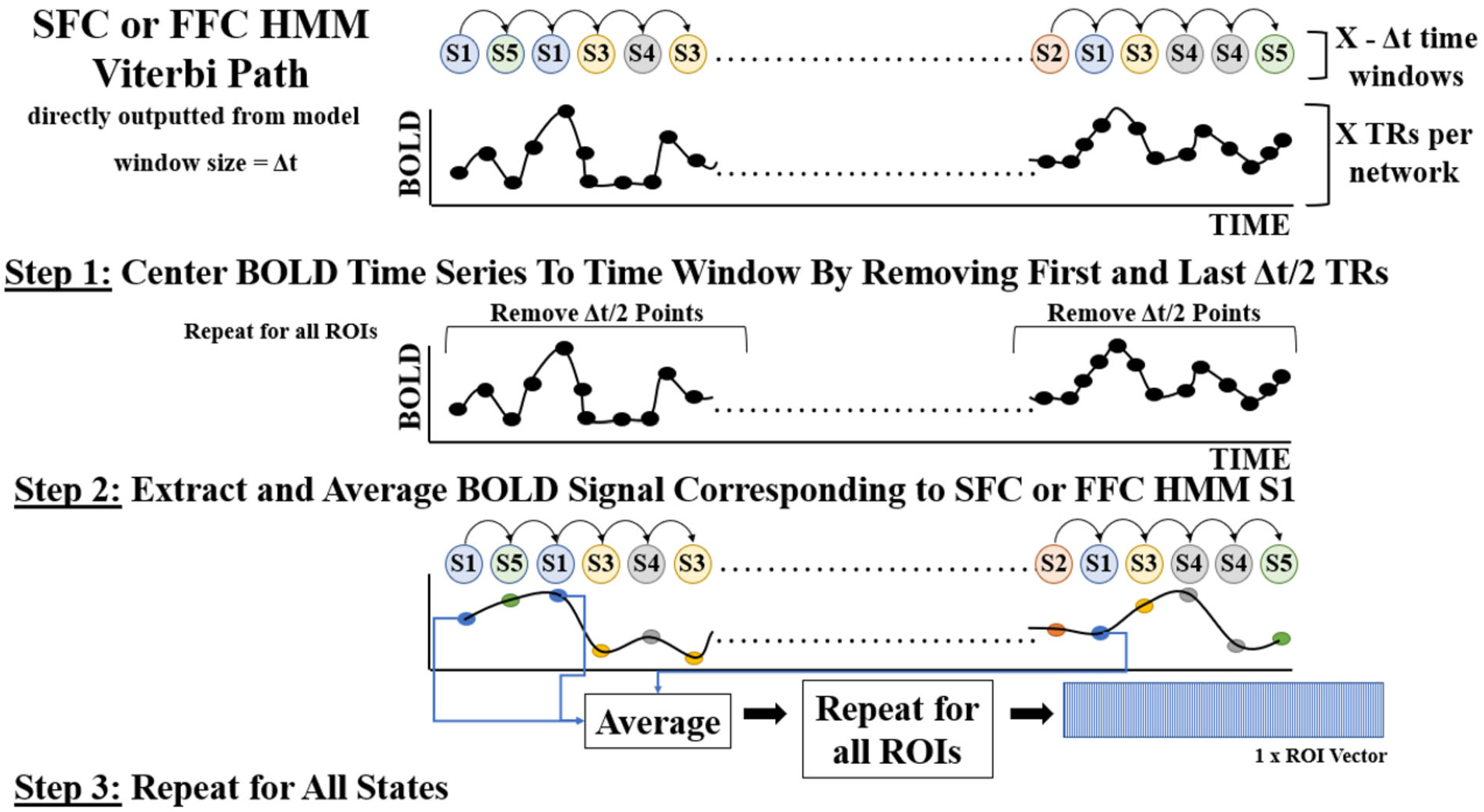
Schematic illustration outlining the method used to recreate the intensity states using the Viterbi paths from SFC and FFC HMMs. The discrepancy in temporal resolution was accounted for by removing some TRs of the BOLD time series to ensure equal length between the BOLD signal and the connectivity-based HMMs’ Viterbi paths.

## B.2 Intensity-based states for all HMMs

The intensity patterns for IB HMM are seen in **Fig. B2** where the subscripts correspond to the model discussed; i.e., S1_IB_ corresponds to state 1 from IB HMM) were directly outputted from the model fitting procedure. S1_IB_ appears to be a DMN-dominant state since the DMN is activated, and all other networks are deactivated. S2_IB_ shows both DMN and FPCN to be activated. S3_IB_ and S4_IB_ both appear to be attention-dominant states: DAN and SN are activated in S3_IB_ while FPCN, DAN, and SN are activated in S4_IB_. S5_IB_ and S7_IB_ both show all networks to be activated, but S5_IB_ has slightly lower activation levels compared to S7_IB_. In S6_IB_, DMN, DAN, and SN are deactivated and FPCN has minor positive activation levels. S8_IB_ shows all networks to be deactivated. The corresponding connectivity states stemming from the output covariance matrices can be found in the bottom row of **Fig. 3**. The intensity states for SFC and FFC HMMs using the methods illustrated in **Fig. B1** are seen in **Fig. B3a** and **B3c**, respectively while the Euclidean distances between them and the intensity states directly outputted from IB HMM are seen in **Fig. B3b** and **Fig. B3d**, respectively. Comparing the two connectivity-based models’ results against IB HMM’s outputted intensity state patterns aimed to determine whether the model types were recognizing different states and ensured that repeated information was not acquired from different HMMs. The similarities of these states to each other were assessed via Euclidean distances (**Fig. B3**).

**Figure B2.**
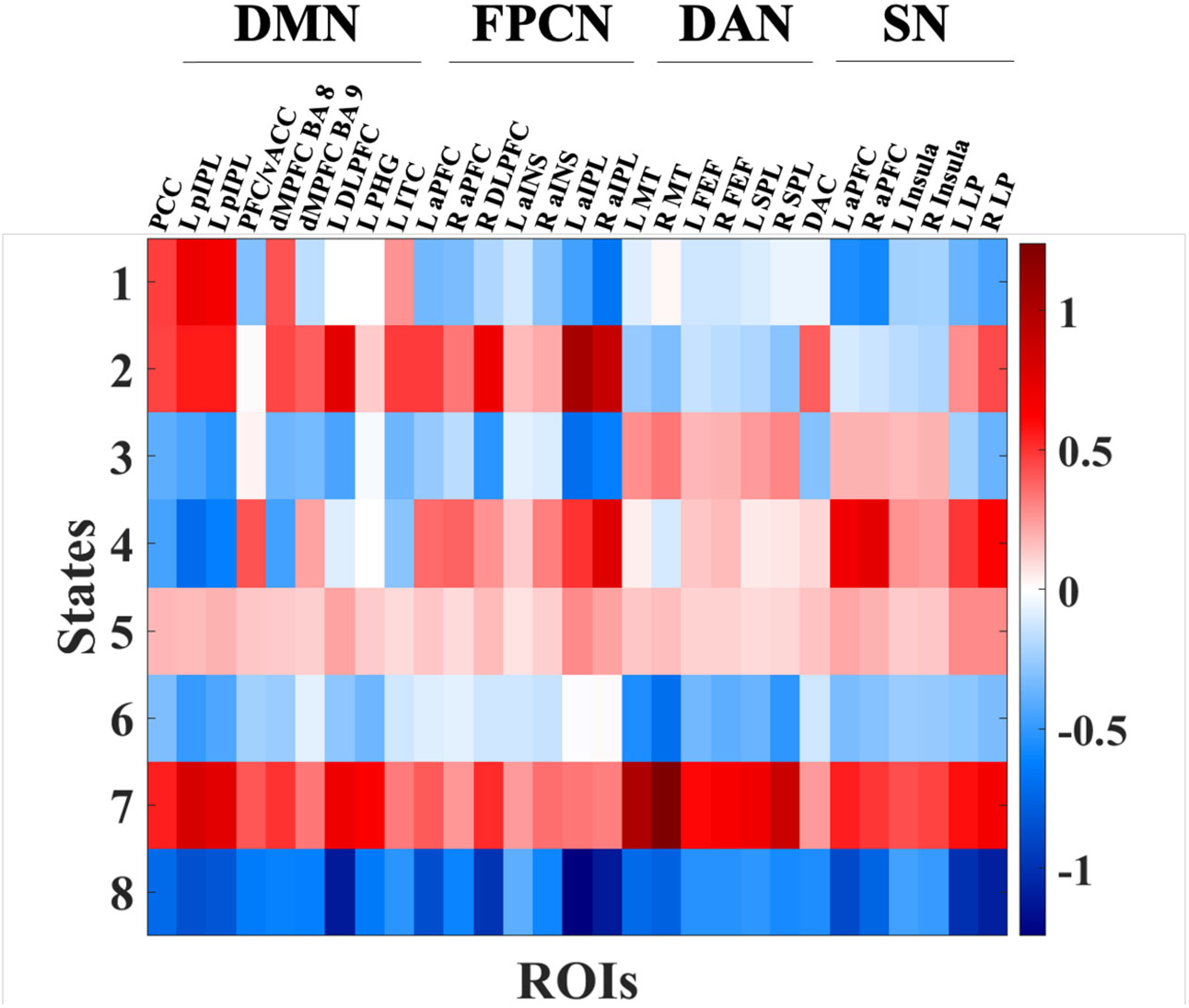
IB HMM activation state patterns. Values shown in red correspond to positive activation levels while those in blue correspond to deactivation.

**Figure B3.**
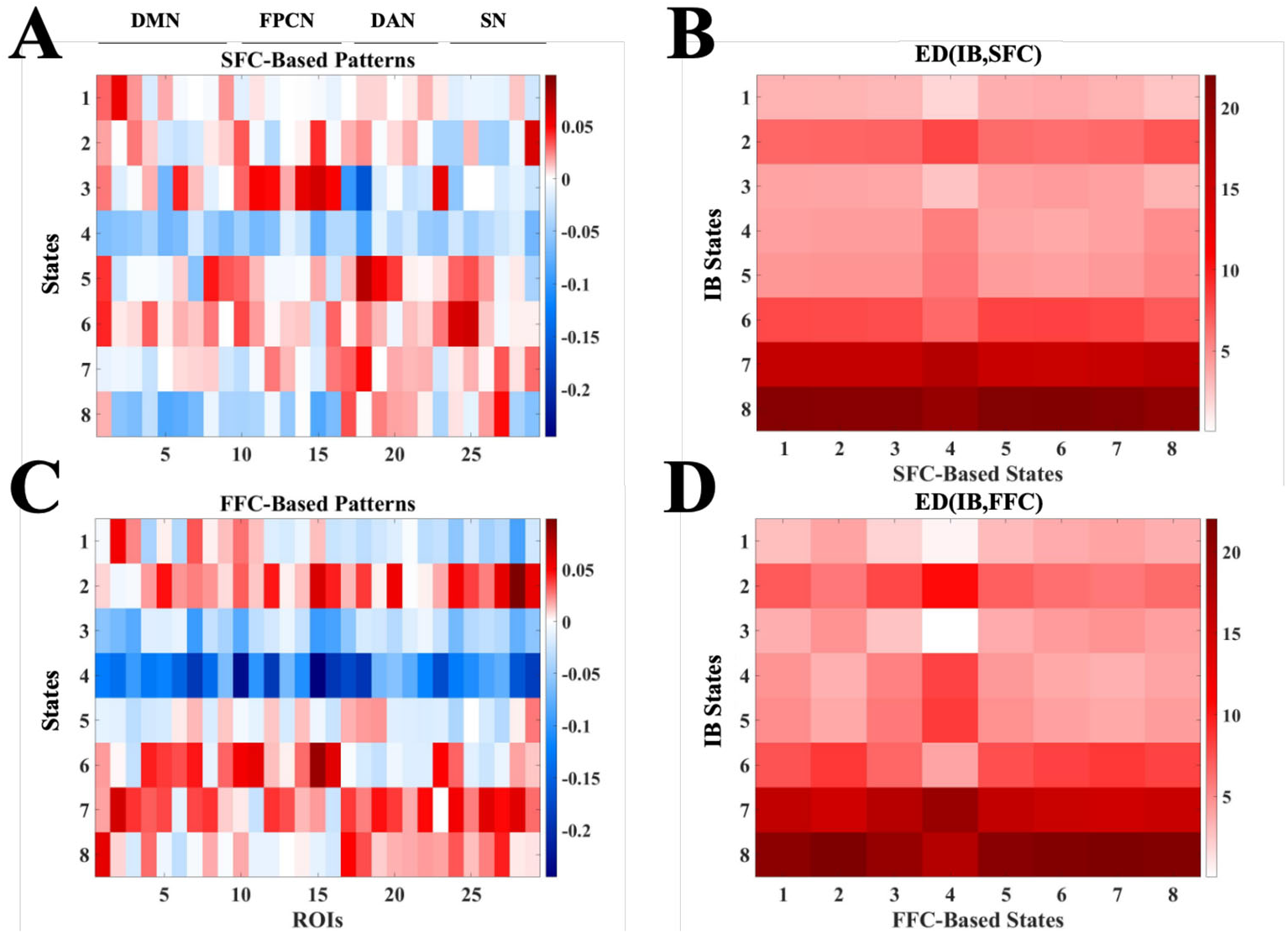
Intensity patterns from averaging the BOLD signal according to the Viterbi path from **(A)** SFC HMM and **(B)** FFC HMM. **(C)** and **(D)**, respectively, show the Euclidean distances between the activation patterns directly outputted from IB HMM (**Fig. B2**) and those from panels **A** and **B**.

These results indicate that IB HMM recognized fluctuations in BOLD signal, and that SFC and FFC HMMs were not sensitive to those same patterns. Each model adequately recognized changes in the data inputted into their respective model type and, therefore, did not recognize the same state patterns. Thus, each HMM is distinct in identifying changes unique to the inputted data, and consequently, in identifying states.

## B.3 Intensity-based state model validation

To create “intensity-defined” ground truth states, eight intensity states (matching the best number of hidden states recovered in the primary analysis; see **Results Section 3.1.1**) were induced by adding an arbitrary value of 2 (the signal is in arbitrary units) to nodes in certain networks and at certain time points within the permuted data.

**Fig. B4a** depicts the time series of states we induced during the validation analysis. In accordance with the stability analysis results, 8 states were induced in our model validation analyses. For the first validation instantiation, intensity states were induced by adding a value of 2 to the specified networks and fitted with IB HMM, which adequately recovered each of the described states (**Fig. B4b**). Compared to the induced state sequence (**Fig. B4a**), the IB HMM fitted to this sequence outputted a state time course with mean R^2^ across all subjects of 0.9999 ± 4.1551e-04. Consistent with the unique nature of intensity-based versus connectivity-based states (see **Results Section 3.2.1**), as expected neither of the connectivity-based models could recover states induced by intensity-based approaches (**Figs. B4c** and **B4d**; SFC HMM R^2^ = 0.0501 ± 0.1003 and FFC HMM R^2^ 0.0739 ± 0.0683 respectively). This finding suggests that our IB HMM model fitting procedure is ideally suited to recover intensity-based states, and the SFC and FFC models do not ‘accidentally’ discover such states when they are present.

**Figure B4.**
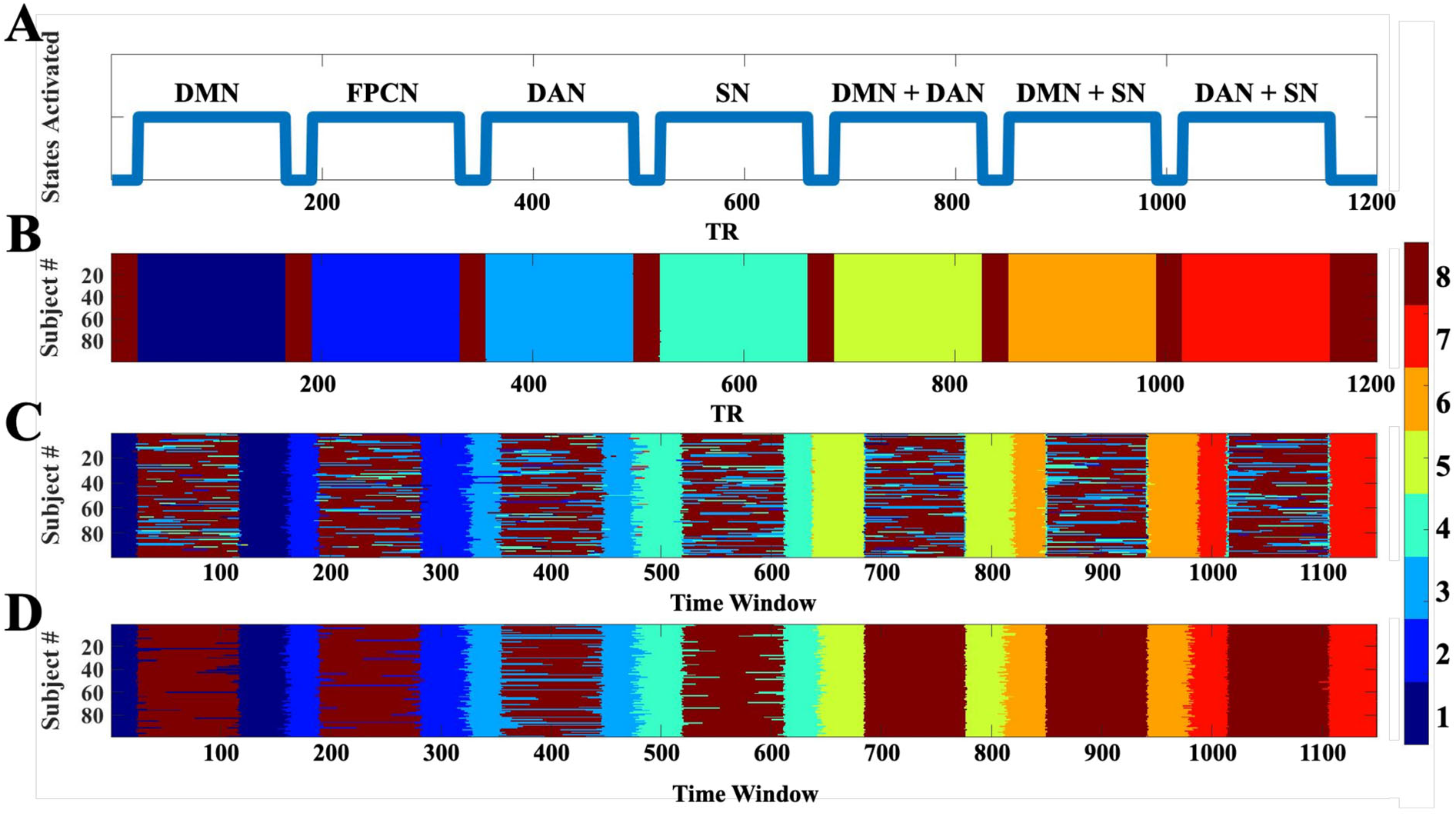
Verification of HMM intensity-based states. **(A)** The artificially induced state sequence depicted which networks exhibited slightly increased network intensity levels. Outputted state sequences from **(B)** IB HMM, **(C)** SFC HMM, and **(D)** FFC HMM when intensity states were induced.

## Appendix C. Additional results

## C.1 Determining the number of hidden states for each model

The RAICAR-based stability analysis results for IB, SFC, and FFC HMMs are shown in **Fig. C1**, limited to models with 6 to 10 hidden states for clarity (full results for models with 3-15 hidden states are shown in **Figs. C2** and **C3** and do not change the conclusions presented in the main text). The smallest number of hidden states at which both IB HMM and SFC HMM demonstrated adequate stability was model order 8 (**Figs. C1a** & **C1b**). We therefore examined the stability of FFC HMM with 8 states, and determined it to be higher than a 9-state model (which had also been relatively stable for both IB HMM and SFC HMM; **Fig. C1c**); thus, in order to demonstrate the clearest and most easily-interpretable comparison among all three HMMs, we assigned FFC HMM to have 8 states as well.

Importantly, setting all models to have the same number of hidden states allowed for a level comparison of local and global analyses across all HMMs, setting FFC HMM on equal footing to the two established models in terms of complexity of outputs. This is important for the purpose of this project, which was a direct comparison among all three models and their relative utility in extracting “pure” connectivity states from resting state fMRI data. However, we note that an identical number of hidden states between any two HMMs may not always occur and users of any of these HMM approaches should not assume that the number of hidden states that is appropriate for one model is appropriate for another. Specifically, there is no mathematical or conceptual basis to suggest that FFC HMM should have the same number of hidden states as SFC HMM or IB HMM especially (nor that SFC HMM and IB HMM should share the same number of states). The fact that we determined 8 states to be best for all models in the present analysis is therefore incidental, and may have occurred because we focused on a relatively small number of ROIs, which greatly narrowed the model complexity.

**Figure C1.**
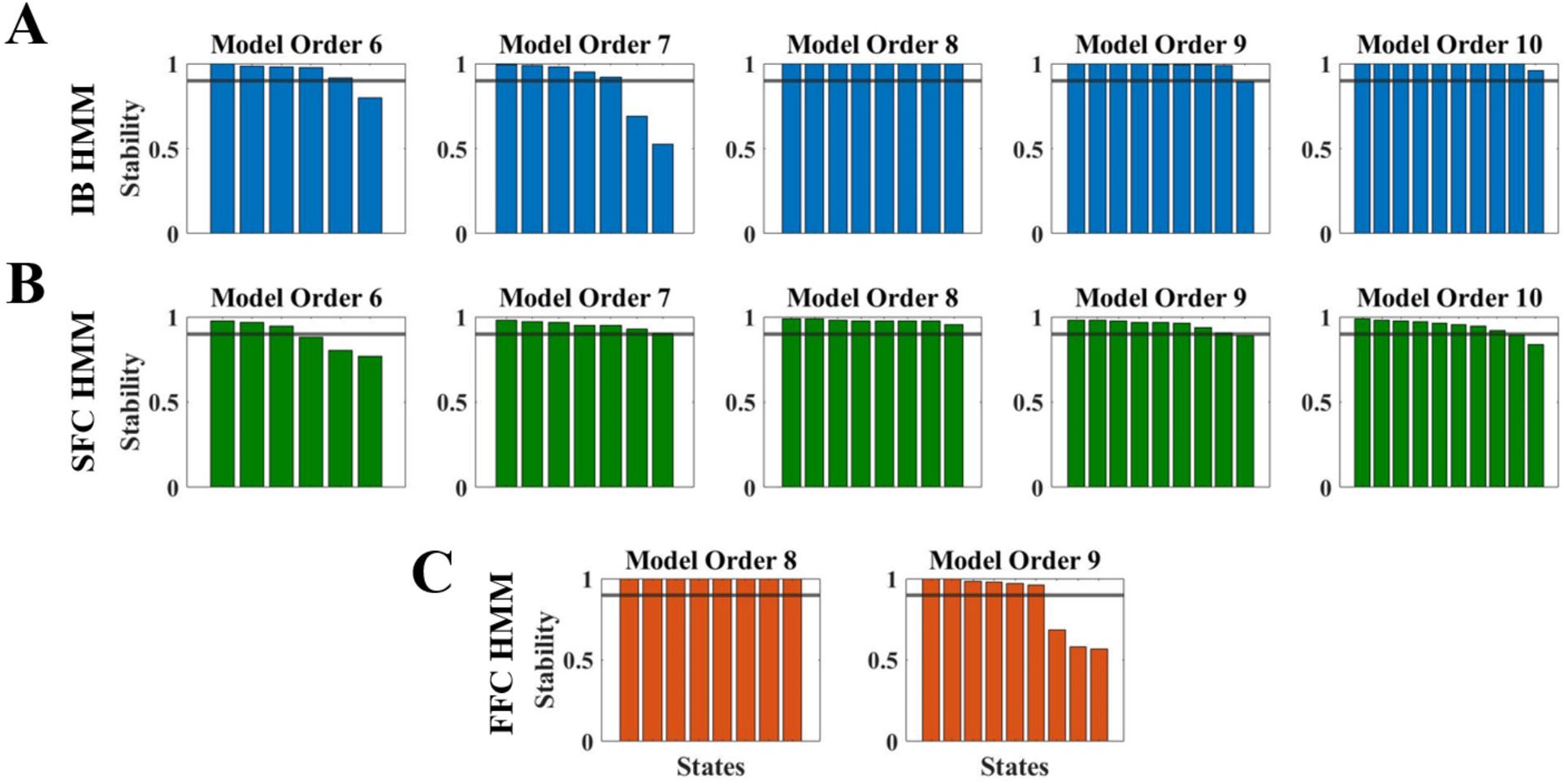
RAICAR-based stability analysis results for **(A)** IB HMM, **(B)** SFC HMM, and **(C)** FFC HMM. Eight hidden states were selected as the best size for all HMMs.

**Figure C2.**
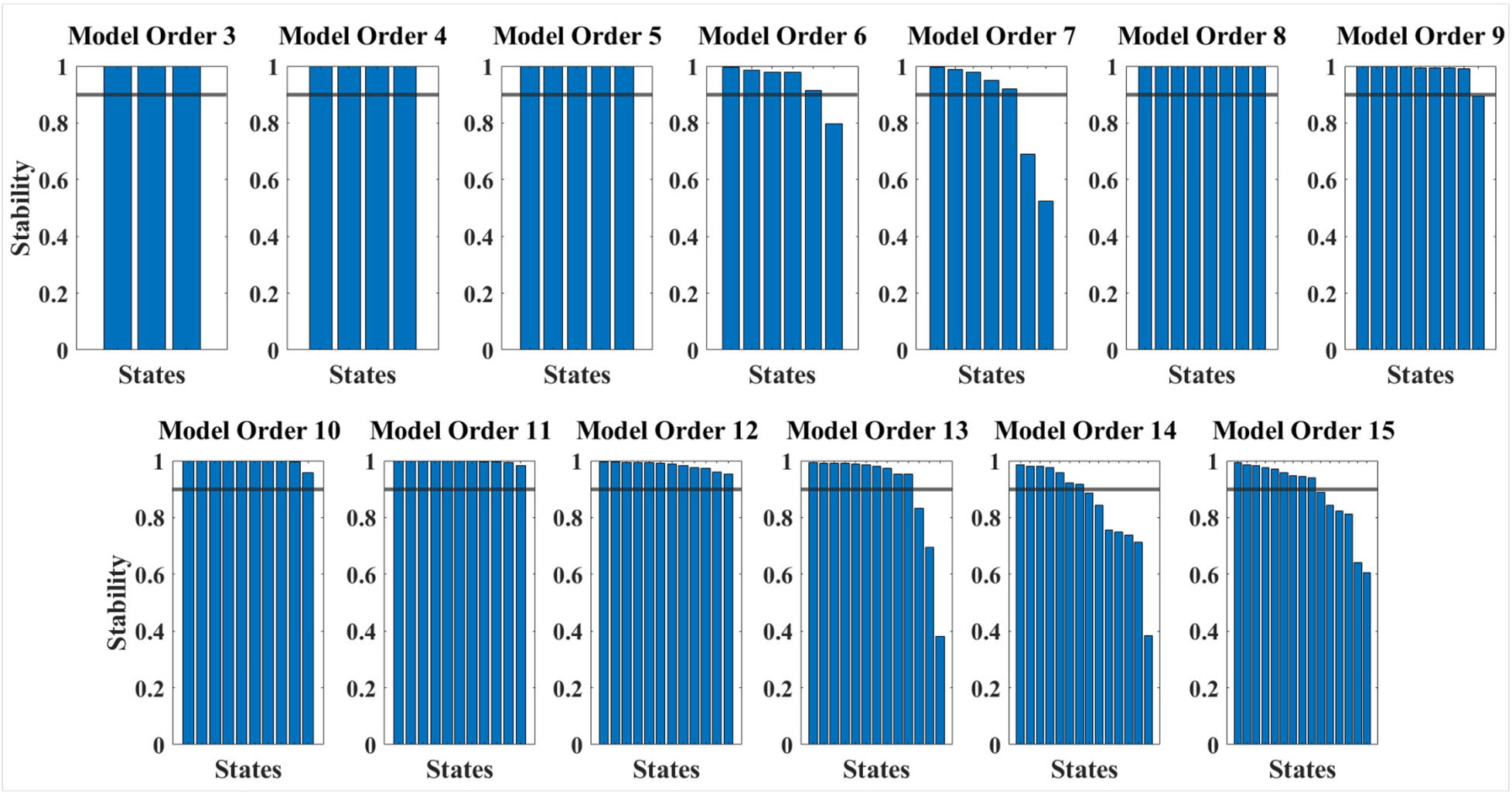
Stability analysis results for IB HMM for models with 3-15 hidden states with a predetermined threshold at 0.9. It may appear that models with as many as 12 hidden states could be suitable for IB HMM because its stability values remained above the predetermined threshold. However, models with 10, 11, and 12 states contained a repeated state making then undesirable as a lack of parsimony likely occurred in the state spatial patterns. The 9-state model identified a state where mean activation equaled zero (consistent with Chen and colleagues’ (2016) findings) that was not observed in the 8-state model (cite once changes are accepted). The 10-, 11-, and 12-state models included two occurrences of this activation pattern; although these models were considered stable, they were undesirable because they contained a repeated state and a lack of parsimony. Thus, eight states were the preferred choice for IB HMM in this investigation.

**Figure C3.**
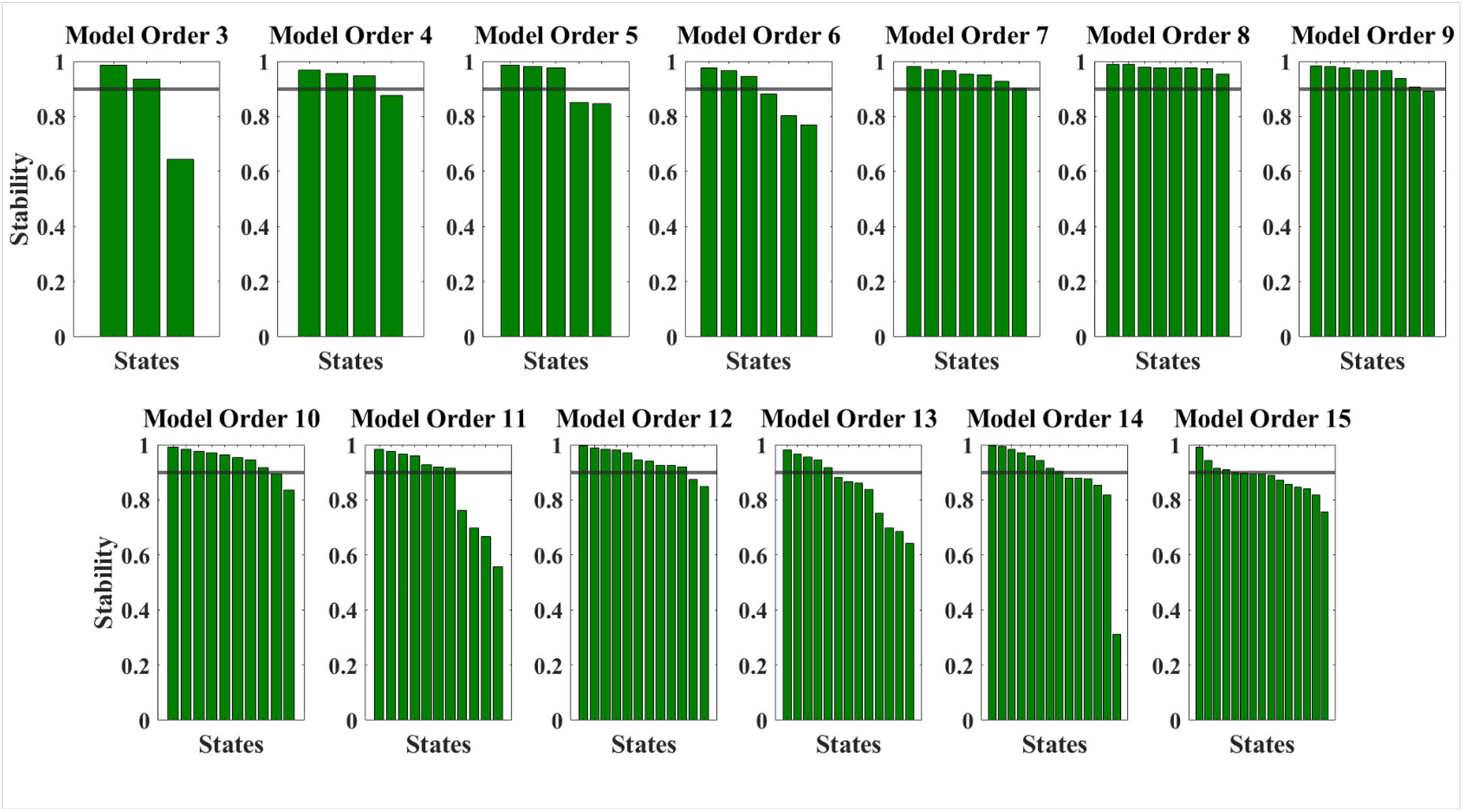
Stability analysis results for SFC HMM for models with 3-15 hidden states with a predetermined threshold at 0.9.

## C.2 Model robustness to data size

Because FFC HMM is a more complex model than either IB HMM or SFC HMM – that is, it must fit more complex full connectivity patterns rather than summed connectivity vectors or intensity-based states – we also need to assess its robustness to the amount of data available to train the model. We also want to compare this to the volume of data necessary to adequately train the comparison models, IB HMM and SFC HMM. This is especially important given that one might wish to use the method with shorter resting state scans or fewer subjects than used here.

To determine the degree to which the states recovered by each of the IB, SFC, and FFC HMMs were dependent on the exact length of the input data, we repeated all fitting procedures on only the first half of the resting state scan (first 7.2 minutes; see **Methods Section 2.3** & **Appendix A.5**). When we halved the dataset, aligned the connectivity states by their similarity, and then computed their overall maximal similarity, we found that the connectivity state patterns largely remained the same for FFC HMM and SFC HMM in particular, but (again unsurprisingly) not for IB HMM (**Fig. C4**). Therefore, it appears that a resting state scan as short as 7.2 minutes may be sufficient to recover the connectivity states discovered by FFC HMM in a scan twice that long.

**Figure C4.**
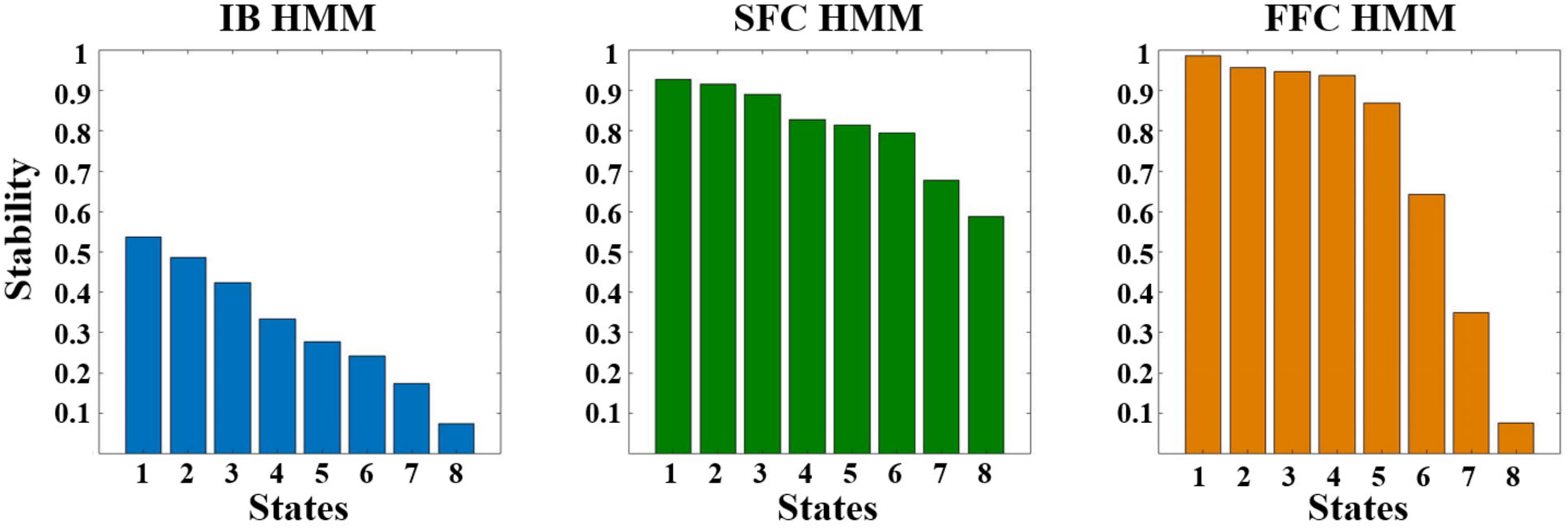
Pearson correlation values (R^2^) between connectivity state patterns identified from whole (14.4 minutes) and half (7.2 minutes) of the HCP Unrelated 100 fMRI dataset for **(A)** IB HMM, **(B)** SFC HMM, and **(C)** FFC HMM.

## C.3 Detailed descriptions of connectivity states

## C.3.1 Intensity-based HMM state descriptions

For IB HMM, DAN in S7_IB_ appeared relatively disconnected from all other networks and associated with activated DMN, FPCN, DAN, and SN. ROIs within DMN and DAN showed slightly above average connectivity in S4_IB_ and were linked to an attention state where FPCN and DAN show the highest levels of activation. Finally, S8_IB_ showed a relative disconnection between ROIs in FPCN and DAN while DAN exhibits slightly elevated within-network connectivity and is paired with all four networks’ deactivation.

## C.3.2 Summed functional connectivity HMM state descriptions

From visual examination, S1_SFC_ showed below-baseline correlations among all networks, particularly within and between DAN and SN, indicating a relative disconnect within the attentional system. Elevated correlations within and between all networks except DAN were observed in S2_SFC_ and DAN did not interact with the rest of the networks within this state. Slightly higher than average correlation values were seen within S3_SFC_ with the exception of several DMN exhibiting below-average connectivity. The strongest connectivity within and between the attentional networks was seen in S4_SFC_. S5_SFC_ had a disconnect among DMN, FPCN, and SN. In S6_SFC_, below baseline connectivity was seen within DAN and between DAN and the other networks; SN exhibited similar behavior. These properties indicate a disconnect within and between these attentional networks. S7_SFC_ showed a slight disconnect between DMN and all other networks indicating a disconnect in the resting state networks. Elevated connectivity within DMN and between DMN and all other networks was seen in S8_SFC_.

## C.3.3 Full functional connectivity HMM state descriptions

S1_FFC_ exhibited no distinguishable state differential functional connectivity states (all connectivity values were around baseline), while S2_FFC_ showed slightly higher than baseline connectivity within and between all networks except for DAN. Above average correlations within and between all networks were seen in S3_FFC_. S4_FFC_ showed all networks to have above average correlations with one another, while S5_FFC_ had slightly above baseline correlations between DAN and all other networks. S6_FFC_ exhibited below baseline correlations between DAN and all other networks as well as between SN and all other networks. DMN was disconnected from all other networks and SN and DAN have slightly above average correlation within and between each other in S7_FFC_. Finally, reduced correlations between DAN and all other networks were seen in S8_FFC_.

## C.4 Time window analysis (connectivity-based HMMs)

One concern might be about whether the use of a sliding window to calculate Pearson correlations before inputting into the HMM violates the Markov property of the HMMs fitted here. That is, HMMs – and all Markov models – explicitly assume that the state occupancy at time t is dependent only on the occupancy at time t-1, i.e. the previously-occupied state, and is independent of previous time periods. Thus, it might be of concern that the use of a sliding window would introduce violations of this Markov property.

To address this concern, we evaluated the outcomes of the fitting procedure as a function of window length used in the sliding-window computation of dynamic functional connectivity. This choice has the potential to greatly impact the states recovered by FFC HMM and SFC HMM (Lurie et al., 2020). Lurie and colleagues postulated that a window length less than 60 seconds may be optimal (Lurie et al., 2020). Thus, for the primary analysis, a window length of 50 time points (36 seconds) was selected. However, to assess the extent to which the qualitative behavior of the model is dependent on window length, we also fit the model under varying window length with sizes of 30 time points (21.6 seconds), 40 time points (28.8 seconds), 50 time points (36 seconds), 60 time points (43.2 seconds), and 80 time points (57.6 seconds). If results are robust to perturbations in this window length beyond predictable increases in switch rate (probability to transition out of a given state at any given time), we can conclude that any violations of the Markov property do not unduly impact the results presented here.

To evaluate the impact of window length used to calculate dynamic functional connectivity on the connectivity states recovered by FFC and SFC HMMs, we therefore examined how connectivity states would change as window length varied from 30 to 80 time points (36-57.6 seconds). Importantly, neither SFC nor FFC HMMs showed a strong effect of such a change in window length. FFC differential functional connectivity states showed minimal differences across window lengths. All states matched to their counterparts with R^2^ ≥ ~0.5 with 1 exception: matching S3_FFC_ 40tp to S3_FFC_ 80tp (R^2^ = 0.3469). Likewise, all SFC differential functional connectivity states matched their counterparts across window sizes with Pearson R^2^ ≥ 0.5 except for two instances: (1) correlating S4_SFC_ 30 time points (tp) to S4_SFC_ 80tp (R^2^ = 0.3084), and (2) S3_SFC_ of any time length to 80tp (tp=30, R^2^ = −0.3587; tp=40, R^2^ = −0.2588; tp=50, R^2^ = −0.3062; tp=60, R^2^ = −0.2563). In fact, although the difference is slight, SFC states may have shown more variability across different time windows than FFC states because of the summing factor, i.e. the definition of connectivity in SFC as the sum of connectivity from one node to all other nodes. As seen in **Figs. C5** and **C6**, poor pattern matching occurred when correlating differential functional connectivity states to differential functional connectivity states from a window size of (80 tp or ~60 seconds).

These results show that a window size containing less than approximately 60 seconds of data is preferred for these two connectivity-based HMMs, consistent with the ideas Lurie et al. 2020 presented (Lurie et al., 2020). The histograms of connectivity values across the differential functional connectivity states (far right columns, **Figs. C5** and **C6**) support this interpretation, since the distributions of connectivity values became less separated as the window size increased; this reduction in distributional separability suggests that the state patterns’ individual distinctiveness -- i.e., lack of resemblance to their counterparts -- was larger in smaller window sizes and was reduced as window length increased. This was particularly noticeable in the 80tp window size, where the distributions of connectivity values appear closest together, meaning the differential functional connectivity states are minimally distinct from one another.

Thus, the window length used in the primary analyses presented above (50 time points, or ~36 seconds) produces similarly unique states as other reasonable window lengths. Together, these analyses confirm that FFC HMM recovers connectivity states that are distinct from those recovered by SFC HMM (and IB HMM), and that the differences between FFC HMM’s connectivity states and those from other models are not trivially due to somewhat arbitrary choices about window length for calculating dynamic functional connectivity measures.

**Figure C5.**
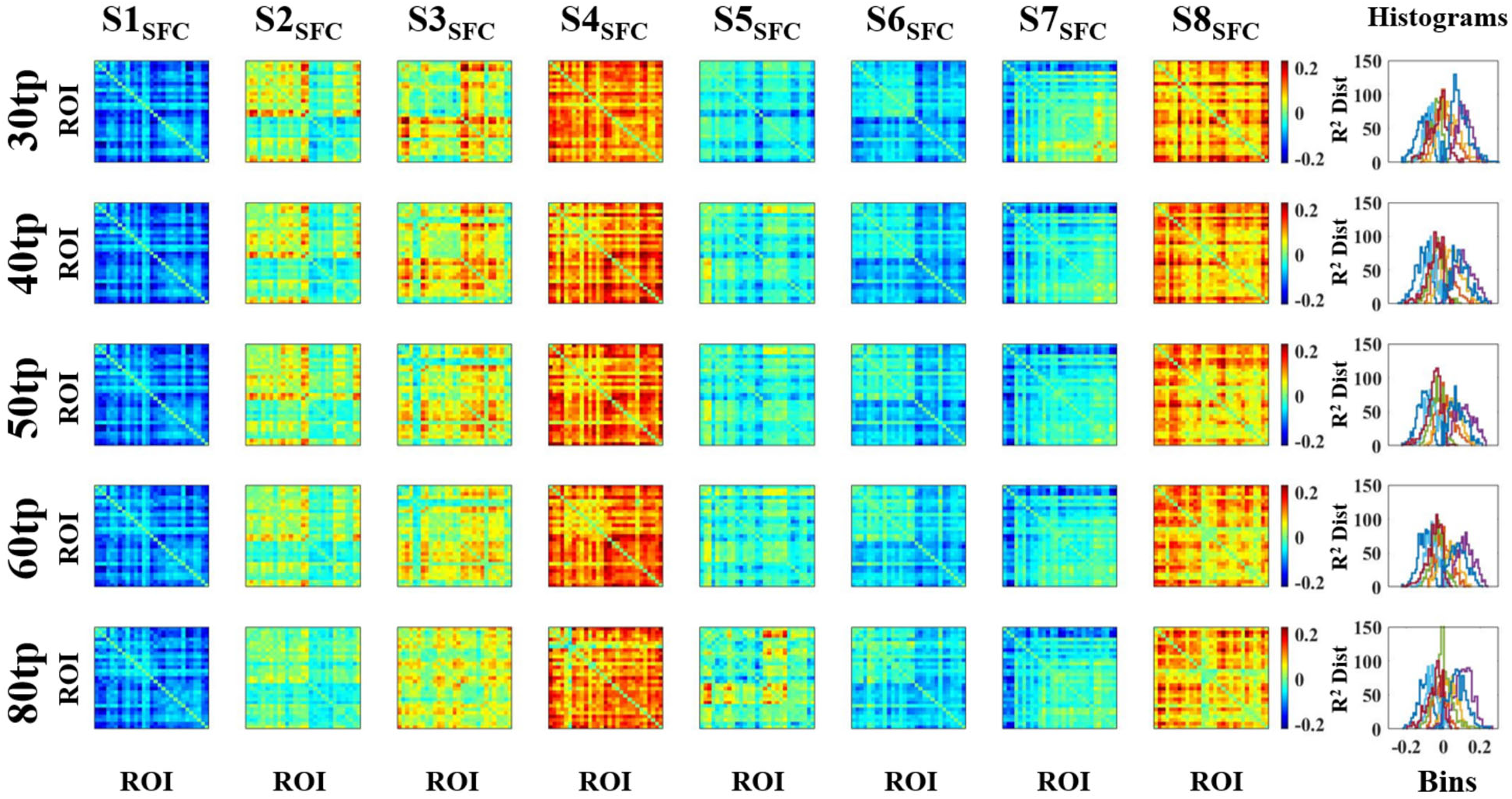
SFC HMM connectivity differential functional connectivity states for window sizes of 30tp, 40tp, 50tp, 60tp, and 80tp used in the sliding window correlation analysis. The histograms (rightmost column) display the overall distribution of R^2^ values of S1_SFC_-S8_SFC_ differential functional connectivity states for each window size. As window size increases, the histograms become less separated, indicating that the states become more similar in pattern as window size increases. These results show that any window size containing less than ~60 seconds of data (60tp or less) produces similarly distinct connectivity profiles for SFC HMM.

**Figure C6.**
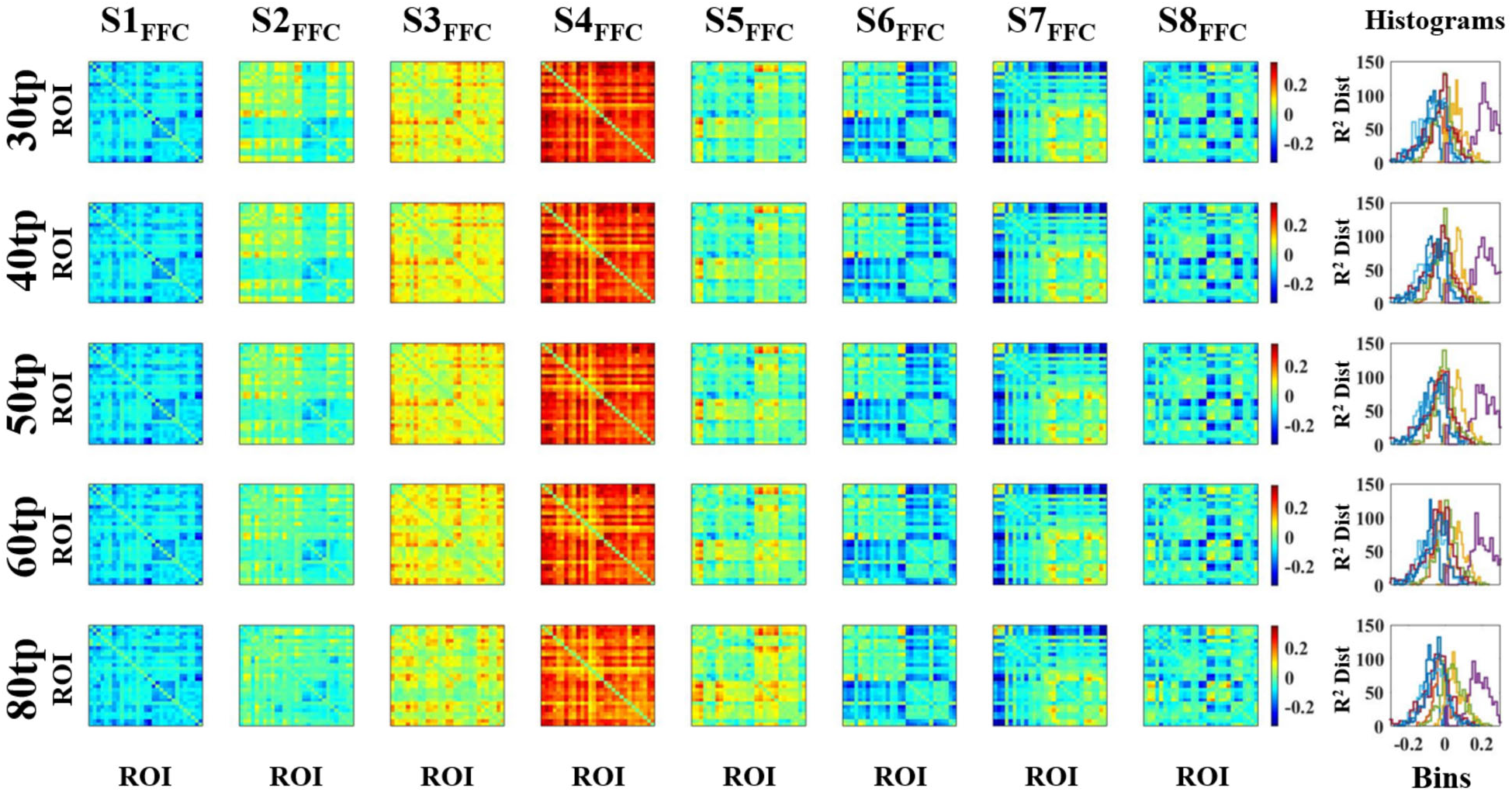
FFC HMM connectivity differential functional connectivity states for window sizes of 30tp, 40tp, 50tp, 60tp, and 80tp used in the sliding window correlation analysis. The histograms (rightmost column) display the overall distribution of R^2^ values of S1_FFC_-S8_FFC_ differential functional connectivity states for each window size. As window size increases, the histograms become less separated, indicating that the states become more similar in pattern as window size increases. These results show that a window size containing less than ~60 seconds of data (60tp or less) produces similarly distinct connectivity profiles for FFC HMM, and that differences between FFC HMM’s connectivity states and those from other models cannot trivially be explained by arbitrary choices about window size.

## C.5 Fisher-z transform does not impact FFC HMM

One possible concern is that FFC HMM may violate assumptions of a Gaussian HMM, i.e. that the off-diagonal elements of the correlation matrices either are distributed normally or that violations of this assumption do not affect results. To test this possibility, we Fisher-z transformed the Pearson correlations outputted by the sliding-window approach for the main results presented here so that they would occupy a range of [−∞,∞] rather than [−1,1], and then re-fit FFC HMM to evaluate any differences in behavior.

This analysis revealed essentially no difference in the recovered functional connectivity states whether or not the Fisher-z transform is applied (**Fig. C7**). Correlations between (matched) states for Fisher-z transformed versus non-transformed inputs showed extremely high and statistically significant consistency, with only one exception which – interestingly – was significantly anti-correlated. Therefore, for the sake of continuity with previously-presented approaches (especially SFC HMM), we opted to present the non-Fisher-z transformed results in the main text.

**Figure C7.**
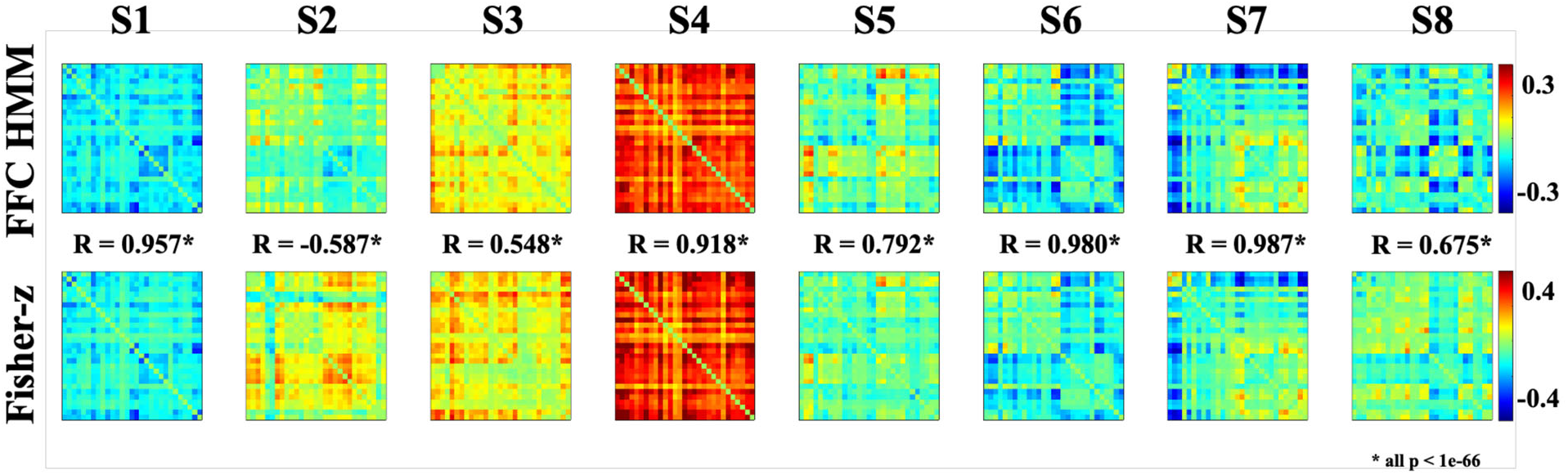
Comparison between FFC HMM across Fisher-z transformed inputs versus raw Pearson correlation inputs. There is a high degree of correspondence between the state differential connectivity profiles recovered by both models, showing that the decision of whether or not to Fisher-z transform the Pearson correlations prior to fitting FFC HMM is unlikely to meaningfully affect the recovered states or their interpretation. The one exception is in S2, which is significantly anti-correlated; however, this significant anti-correlation may be due to the reliance here on state highlights rather than connectivity states themselves.

